# Spontaneous network activity links misrouted interneuron migration to cortical maturation abnormalities

**DOI:** 10.64898/2026.05.07.723521

**Authors:** Sathish Venkataramanappa, Jürgen Graf, Holger Haselmann, Dagmar Schütz, Oliver Storch, Dominic Römer, Praveen Ashok-Kumar, Nelli Blank-Stein, Christian Mayer, Knut Holthoff, Knut Kirmse, Philipp Abe, Ralf Stumm

**Affiliations:** Institute of Pharmacology and Toxicology, Jena University Hospital, 07747 Jena, Germany; Department of Neurology, Jena University Hospital, 07747 Jena, Germany; Max Planck Institute of Neurobiology, 82152 Martinsried, Germany; Department of Neurophysiology, Institute of Physiology, University of Würzburg, 97070 Würzburg, Germany; Institute for Clinical Genetics, University Hospital Carl Gustav Carus at TUD Dresden University of Technology and Faculty of Medicine of TUD Dresden University of Technology, 01307 Dresden, Germany; ThIMEDOP – CeTraMed, Jena University Hospital, Friedrich Schiller University Jena, Am Klinikum 1, 07747 Jena, Germany; Developmental Biology of the Immune System, Life & Medical Sciences (LIMES) Institute, University of Bonn; 53115 Bonn, Germany

## Abstract

Cortical inhibitory neurons (CIN) populate the neocortex and hippocampus by extensive tangential migration. This process is highly vulnerable to genetic and environmental disturbances and is linked to neuropsychiatric disorders. However, the mechanisms by which transient migratory abnormalities translate into persistent functional deficits remain poorly understood. Here, we utilized a conditional *Cxcr4* knockout to investigate the consequences of disrupted migratory guidance - a feature of genetic schizophrenia models. This demonstrated that despite migrating in ectopic cortical layers, CIN quantitatively colonized the neocortex until birth, whereas limbic regions developed a permanent deficit in CIN numbers. Furthermore, CIN failed to populate the neocortical marginal zone, a transient reservoir for late-born CIN destined for superficial cortical layers. Consequently, the layering and molecular identities of CIN, as well as the synaptic connectivity and spontaneous activity of the neuronal network, were significantly altered in the early postnatal neocortex. Although abnormal CIN layering was gradually compensated before maturity, functional differences persisted, as evidenced by facilitated propagation of sensory stimuli between cortical areas. These results demonstrate that CIN migration is an instructive process critical for activity-driven early postnatal network maturation and CIN identity, thus providing a mechanistic link between migratory guidance and the integrity of mature circuits.

## Introduction

CIN play a critical role in shaping the maturation of the cerebral cortex at various stages of development (1, 2). As CIN originate in the ventral telencephalon, they migrate considerable distances to reach their destinations (3). This process unfolds over an extended developmental period and, when affected by genetic or external disturbances, may contribute to the development of psychiatric disorders (4–8). Notably, many of these clinical conditions lack overt numerical deficits or gross laminar disorganization of CIN (4), suggesting that pathogenic mechanisms may instead emerge from perturbed CIN maturation and altered circuit formation. We hypothesized that migration is not merely a transport process, but also an instructive mechanism through which CIN acquire anatomical and functional properties necessary in the early postnatal period, during which CIN regulate activity-dependent refinement of the developing cortical network (1, 2).

The majority of CIN derive from NKX2.1^+^ progenitors in the medial ganglionic eminence (mCIN). These comprise Somatostatin^+^ (SST), Parvalbumin^+^ (PVALB), and a fraction of LAMP5^+^ neurons (9, 10). In mice, mCIN invade the cortex by embryonic day E12.5, enter the cortical marginal zone (MZ) and subventricular-intermediate zone, and undergo tangential migration (TM) in these layers to colonize the neocortex and hippocampus until birth (11–16).

TM relies on an intricate interplay of guidance factors (17), with CXCR4 generating the dominant attractant signal for the MZ and subventricular-intermediate zone (13, 18–25). The critical role of CXCR4 in CIN development is underscored by findings linking a defective CXCR4 pathway in CIN to brain disorders including schizophrenia (26–31).

Still, little is known how defective guidance of tangentially migrating CIN may translate into permanent functional deficits. Since mCIN identity is dynamically acquired during migration (32), cues from an abnormal environment could change mCIN maturation and function. Furthermore, proper early guidance might be relevant for the final layering of mCIN. This, however, remains controversial, because different results were obtained with transplantation of *Cxcr4*^-/-^ mCIN (13, 33) and conditional *Cxcr4* deletion strategies (24, 34).

Crucially, CXCR4 is down-regulated in mCIN during the perinatal period, when mCIN switch from TM to radial migration to reach their final laminar position (11, 18, 35–37). This restricts CXCR4 function in mCIN mainly to the TM period, making a *Cxcr4* knockout in mCIN (*Cxcr4*-cKO) an attractive model for studying functional consequences of perturbed mCIN guidance. To achieve this, we utilized *Nkx2.1-*Cre (9) in combination with the *Cre-*reporter Ai14, allowing us to trace and manipulate the mCIN lineage from the progenitor stage onward. Using this model, we address several key questions with respect to guided mCIN migration. These include its influence on the postnatal maturation, molecular identity, and final layering of mCIN, as well as its influence on the architecture and activity of the forming cortical network. Our findings highlight a critical link between migratory guidance and functional maturation of the early postnatal cortical network.

## Results

### Misrouted TM disrupts mCIN layering in the neocortex at birth

In this study, we employed a conditional *Cxcr4* knockout (*Cxcr4*-cKO) to investigate how disruption of the layer-specific migration of mCIN in the embryonic cortex influences the functional assembly of cortical circuits. We identified cells undergoing Cre-mediated genetic recombination via the tdTomato (tdT) reporter signal of the Ai14 reporter line. In the neocortex of *Cxcr4*-cKOs, these cells lacked CXCR4 expression at embryonic stages E13.5 and E16.5 (demonstrated for E16.5 in **Figure S1A**) and were redistributed from MZ and subventricular-intermediate zone towards the cortical plate **(Figures S1B - S1D)**. This defect was comparable to the mCIN positioning defect observed in *Cxcr4*^-/-^ mice (36). Collectively, these findings indicate efficient *Cxcr4* deletion in *Cxcr4*-cKOs. Unlike *Cxcr4*^-/-^ mice, which die prenatally, *Cxcr4*-cKOs were viable postnatally, exhibiting normal body weight and survival **(Table S1)**. Therefore, the *Cxcr4*-cKO model enabled us to bypass early developmental mortality and evaluate the specific consequences of mCIN misrouting at postnatal stages.

Studies using explant technologies led to the concept that CXCR4-signaling increases forward migration of mCIN to promote effective colonization of the cortex (14, 20, 38–40). Since this hypothesis has not been systematically tested in *in vivo* models, we quantified the mCIN density in *Cxcr4*-cKOs across neocortical and hippocampal regions at postnatal day zero (P0), which corresponds to the endpoint of the TM period. We used two independent approaches to identify mCIN: *in situ* hybridization for the mCIN marker *Lhx6*^+^ and immunofluorescence labeling of PDGFRA^-^ tdT^+^ cells **(Figures S2A** and **S2B)**. The early oligodendrocyte marker PDGFRA was used in studies until P8, because *Nkx2.1*-Cre-labeled cells include some first-wave oligodendrocytes that vanish during the first postnatal week (9, 41). We assessed rostro-caudal dispersion in four sectional planes along the rostro-caudal axis by placing a parasagittal region of interest (ROI) in the motor cortex rostrally and the visual cortex caudally. Surprisingly, this revealed no major differences between controls and *Cxcr4*-cKOs (**Figures 1A** and **S2C**). Latero-medial dispersion, analyzed in a rostral and a caudal plane, was affected, as evidenced by reduced mCIN densities in the anterior cingulate area and the hippocampus (**Figures 1B** and **S2C** - **S2E**), regions frequently implicated in neuropsychiatric disorders.

**Figure 1.**
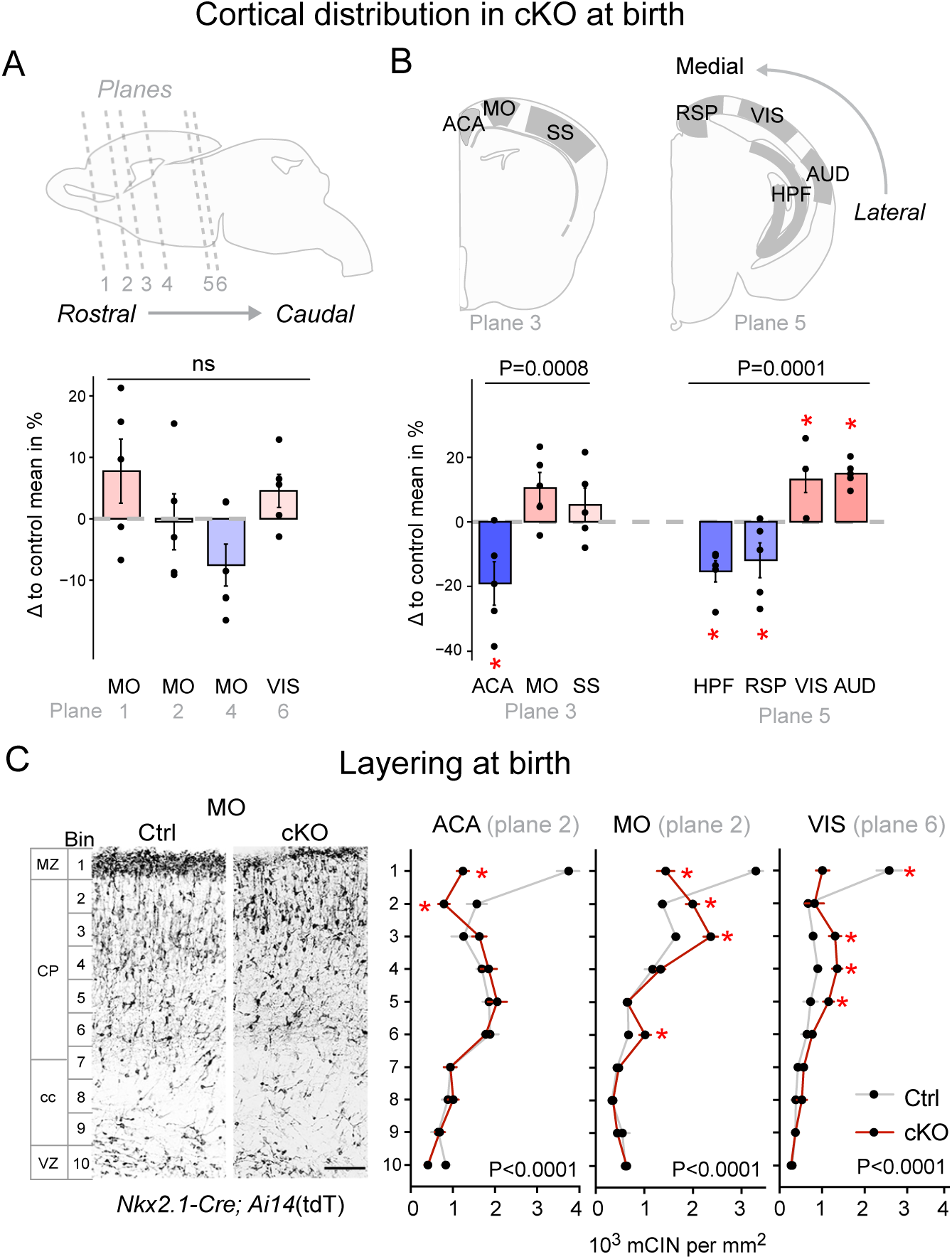
Abnormal regional distribution and layering of mCIN in P0 *Cxcr4*-cKO mice. **A,B, Top,** Schematics illustrate the position of the analyzed coronal sectional planes and cortical areas in the P0 mouse brain. **Bottom**, Deviation in the mCIN density in cortical areas of *Cxcr4*-cKO mice along the rostro-caudal (left) and medio-lateral (right) axes expressed as difference between *Cxcr4*-cKO and control means in percent of the control mean. Quantification is derived from countings of PDGFRA^-^tdT^+^ (***A***, *Nkx2.1*-Cre; Ai14:*R26*^LSL-tdT^) and *Lhx6*^+^ (***B***) cells. Color code indicates the relative abundance of mCIN in *Cxcr4*-cKO (blue: less abundant; red: more abundant). **C**, Micrographs demonstrate tdT in the motor cortex (sectional plane 2). Line graphs show the distribution of mCIN (PDGFRA^-^ tdT^+^) in the indicated cortical regions. mCIN were assigned to bins according to their relative position between pia mater and ventricle. **Statistics: A,B,** Analyses were performed using absolute cell densities as shown in **Figures S2C** and **S2D**. **C,** p-values for bin-genotype interactions as by two-way ANOVA with repeated measures; *: false discovery rate (q) <0.1 as by two-stage linear step-up procedure. **A-C,** Data are mean +/- SEM. **Cortical regions:** AUD, auditory; ACA, anterior cingulate; HPF, hippocampus formation; MO, motor; RSP, retrosplenial; SS, somatosensory; VIS, visual. **Abbreviations:** cc, corpus callosum; cKO, *Cxcr4*-cKO; CP, cortical plate; Ctrl, control; MZ, marginal zone; VZ ventricular zone. **Scale bar:** 100 µm in ***C.*** Data are available in **Table S5**.

Next, we examined the layering of mCIN across neocortical regions (**Figures 1C, S2F,** and **S2G),** dividing the cortex into ten bins ranging from the MZ (bin #1) to the ventricular zone (bin #10). In all regions examined, the MZ consistently showed the highest mCIN density in controls and a strongly reduced mCIN density in *Cxcr4*-cKOs. In all regions except in the anterior cingulate area, this reduction was accompanied by an increased mCIN density in the layers subjacent to the MZ. Thus, while CXCR4-guidance of TM is not strictly required for mCIN to effectively colonize the neocortex, it is indispensable for their laminar allocation at birth. Furthermore, less mCIN reached medial limbic structures including the hippocampus.

### The MZ as an intermediate station for mCIN populating late-forming superficial neocortical layers

After completing TM by the time of birth, mCIN initiate radial migration to attain their final laminar position during the first postnatal week. We examined this process and how it is altered when mCIN are forced to migrate radially from ectopic deep-layer locations in *Cxcr4*-cKOs. To this end, we assessed mCIN stratification in layers 1 through 6 of the motor cortex at multiple time points between P0 and P8. We visualized the time- and genotype-dependent alterations in mCIN lamination by depicting the relative mCIN densities for ten cortical bins as heatmaps (**Figures 2A** – **2C;** for line-graphs, see **Figure S3A**). Immunostainings for TBR1 and CTIP2 were performed to define superficial layers (SL: TBR1^-^CTIP2^-^) and deep layers (DL: CTIP2^+^ or TBR1^+^), respectively (**Figure 2B)**. Both, control and *Cxcr4*-cKOs, showed a declining mCIN density in bin #1, though the decline was less pronounced in mutants due to their already low mCIN density in bin #1 at P0. Strikingly, while the mCIN density in bin #2 increased over time in controls, it decreased in *Cxcr4*-cKOs **(Figure 2C)**, indicating a failure of mCIN to populate the layers beneath layer 1. This became more apparent by plotting the fold change in *Cxcr4*-cKOs (**Figure 2C**). Direct comparison of positioning defects at P0 and P8, quantified as absolute mCIN densities, further confirmed these findings (**Figures S3C** and **S3D)**. When assessing the morphology of mCIN during the first postnatal week, we noticed mCIN with a downward-directed migratory morphology beneath layer 1 up to P4 (**Figure 2D**). These data suggest that the MZ serves as a reservoir for mCIN populating late-forming superficial layers. In *Cxcr4*-cKOs, the failure to populate this reservoir during the embryonic period precludes the normal radial redistribution of mCIN, resulting in a numerical shortage in superficial layers.

**Figure 2.**
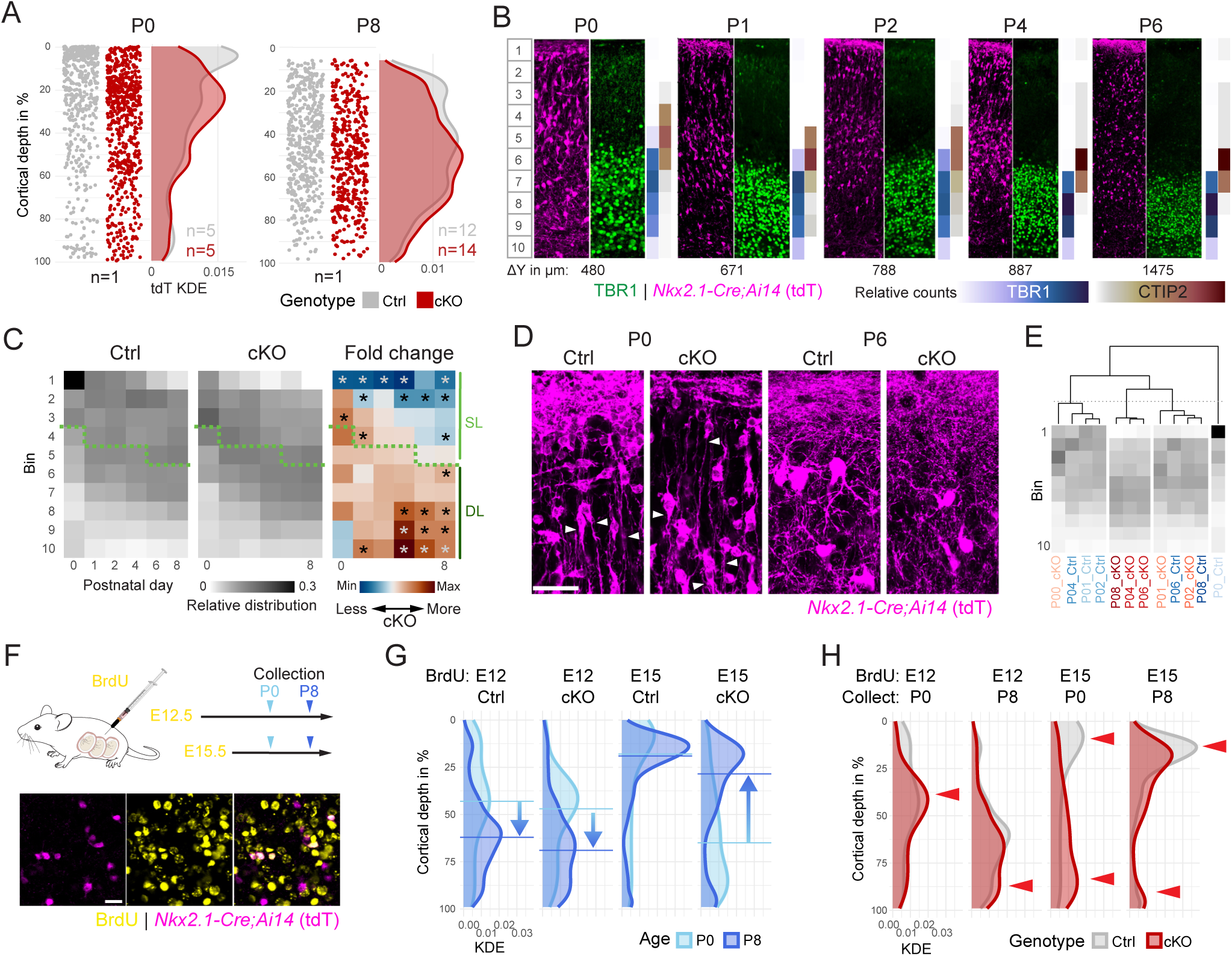
Perturbed TM influences postnatal laminar allocation of mCIN. **A**, Dot plots and curves show the distribution of mCIN across layers 1 to 6 of the motor cortex at P0 and P8. The position within the cortex is represented as relative cortical depth on the y-axis. Dot plots depict individual mCIN. Curves show kernel density estimates (KDE) on the x-axis. n-numbers represent individual mice. **B,** Confocal images demonstrate double immunofluorescences for TBR1 and tdT in layers 1 to 6 of the motor cortex of control mice. Cortical bins are shown to the left, cortical thickness is given as deltaY. Color codes represent cell counts of TBR1 and CTIP2 per bin in percent of the total counted cells. **C**, Heatmaps show the relative density of mCIN across bins #1 to #10 at the indicated postnatal days for control and *Cxcr4*-cKO mice and as the fold change of *Cxcr4*-cKOs versus controls. *p <0.1, beta regression followed by simultaneous tests for general linear hypotheses for repeated measurements correction. Data in ***C*** refer also to **Figure S3A**. Green dotted lines segregate superficial and deep layers. **D**, Z-projections of confocal images demonstrate tdT in layer 1 and subjacent layers of the motor cortex. Arrowheads point to mCIN exhibiting a downward directed migratory morphology. **E,** Unbiased clustering of the mCIN lamination patterns at indicated time points and conditions. Data in ***E*** refer summarize data shown in **Figure S3B**. **A,C,E,** Data represent PDGFR^-^ tdT^+^ cells (i.e. mCIN). **F,** Schematic illustrating the BrdU pulse-labeling paradigm used in ***G*** and ***H***. Confocal images demonstrate a representative double immunofluorescence for tdT and BrdU in layer 2 at P8. **G,H,** Density plots show the distribution of BrdU^+^ tdT^+^ cells (mCIN including some first-wave oligodendrocytes) across layers 1 to 6 with the relative cortical depth on the y-axis and KDE on the x-axis. Horizontal lines show the median. Arrows in ***G*** highlight the shift of the median of the pulsed cohorts between P0 and P8. Arrowheads in ***H*** identify cortical segments with significant frequency differences between control and *Cxcr4*-cKOs. Data in ***G*** and ***H*** refer to **Figure S3F**. Source data are provided in **Table S5**. **Abbreviations:** cKO: *Cxcr4*-cKO; Ctrl, control; DL and SL, deep and superficial cortical layers; KDE, kernel density estimates. **Scale bars:** 33 µm in ***D***, 20 µm in ***F***.

Hierarchical clustering of the mCIN lamination patterns showed that P0 *Cxcr4*-cKOs clustered with P1 - P4 controls and P1 - P2 *Cxcr4*-cKOs clustered with P6 - P8 controls, indicating premature mCIN lamination in *Cxcr4*-cKOs at early postnatal stages. By contrast, patterns of *Cxcr4*-cKOs from P4 to P8 did not cluster with any of the control groups, indicating a persistent lamination defect (**Figures 2E** and **S3B**).

Like principal neurons, mCIN undergo age-dependent stratification, with late-born mCIN colonizing superficial and early-born mCIN colonizing deep layers. This is achieved through radial mCIN migration (11, 42). We investigated how this is affected in *Cxcr4*-cKOs using BrdU pulse-labeling, injecting BrdU at E12 or E15 and analyzing cell layering at P0 or P8 **(Figures 2F – 2H, S3E**, and **S3F)**. Because BrdU immunohistochemistry interferes with PDGFRA detection, we omitted PDGFRA staining in these studies (based on tdT/ PDGFRA double-stainings, <3.6% of the tdT^+^ cells were oligodendrocytes at P0 and virtually none were oligodendrocytes at P8 in controls and *Cxcr4*-cKOs; we therefore refer to E12-pulsed tdT^+^ cells as mCIN).

First, we quantified the layer-specific distribution of total BrdU^+^ cells to assess postnatal layer-formation in *Cxcr4*-cKOs. In controls, E12-pulsed cells populated mainly deep layers at P0 and P8 (Figure **S3E:** 1^st^ diagram), whereas E15-pulsed cells shifted from deep to superficial layers between P0 and P8 (Figure **S3E:** micrographs and 3^rd^ diagram). The latter finding indicates that the superficial layers beneath layer 1 are established, in part, by late-born cells arriving during the first postnatal week as described (42). These processes were similar in control and *Cxcr4*-cKO mice **(Figure S3E)**.

Next, we turned to mCIN. Early-born mCIN (E12 pulse) shifted towards deep layers between P0 and P8 in both genotypes (**Figure 2G**: 1^st^ and 2^nd^ diagram). Direct comparison of control and *Cxcr4*-cKOs showed that early-born mCIN were increased in the middle of the cortex at P0 and in deep layers at P8 (**Figures 2H** and **S3F:** 1^st^ and 2^nd^ diagrams). Late-born mCIN colonized mainly superficial layers in both genotypes at P8 (**Figure 2G**: dark blue in 3^rd^ and 4^th^ diagram). However, the origins of the recruited cells differed: while the superficial layers of controls were populated by mCIN migrating from the MZ and the layers subjacent to the MZ, the superficial layers of *Cxcr4*-cKOs were populated mainly by mCIN migrating from deep layers (**Figure 2G**: light blue in 3^rd^ and 4^th^ diagram). This difference is attributed to the aberrant deep positioning of E15-pulsed mCIN in P0 *Cxcr4*-cKOs (**Figures 2H** and **S3F:** 3^rd^ diagrams). Critically, in *Cxcr4*-cKOs, a significantly larger fraction of late-born mCIN remained in deep layers until P8, leading to a persistent reduction of these cells in superficial layers (**Figures 2H** and **S3F**: 4^th^ diagrams).

Our BrdU labeling experiment shows that attraction of late-born mCIN to postnatally forming superficial layers and of early-born mCIN to deep layers is largely preserved in *Cxcr4*-cKOs. This process partially compensates the gross initial mCIN lamination defect in these mutants. However, a significant fraction of late-born mCIN remained misplacement in deep layers. Consequently, *Cxcr4*-cKOs developed an mCIN deficiency in superficial layers by P8. In sum, TM is critical for mCIN to populate the perinatal MZ which serves as an intermediate station, from where mCIN subsequently populate late-forming superficial layers by radial migration.

### Altered maturation and molecular identity of mCIN at early postnatal stages

Absence of CXCR4 signaling and mispositioning could potentially influence the mCIN transcriptome and identity. We thus conducted bulk RNA-sequencing on sorted E16.5 mCIN (PDGFRA^-^ tdT^+^ cells) (**Figures S4A** and **S4B**). Surprisingly, neither principal component analysis nor Pearson correlation followed by hierarchical clustering revealed clear differences between genotypes (**Figures S4C** and **S4D**). While *Cxcr4* stood out as differentially expressed gene (DEG), only 7 additional DEGs were detected (**Figure S4E),** suggesting that neither CXCR4 signaling nor the local environment significantly shaped the gene expression profile of mCIN by E16.5.

Next, we performed single-cell RNA sequencing (scRNA-seq) on sorted mCIN at P6 (**Figures 3A** and **S5A**), which corresponds to the end of the active migration period. This approach allowed us to identify mCIN subtypes through transcriptomic profiling (32), bypassing the limitations of immunohistochemistry at early postnatal stages, when markers such as PVALB have not yet reached detectable protein levels. UMAP visualization showed a similar global distribution of mCIN from controls and *Cxcr4*-cKOs within the embedding space. However, regional differences in cellular density were apparent (**Figures 3B**), suggesting altered molecular profiles and potential identity shifts.

**Figure 3.**
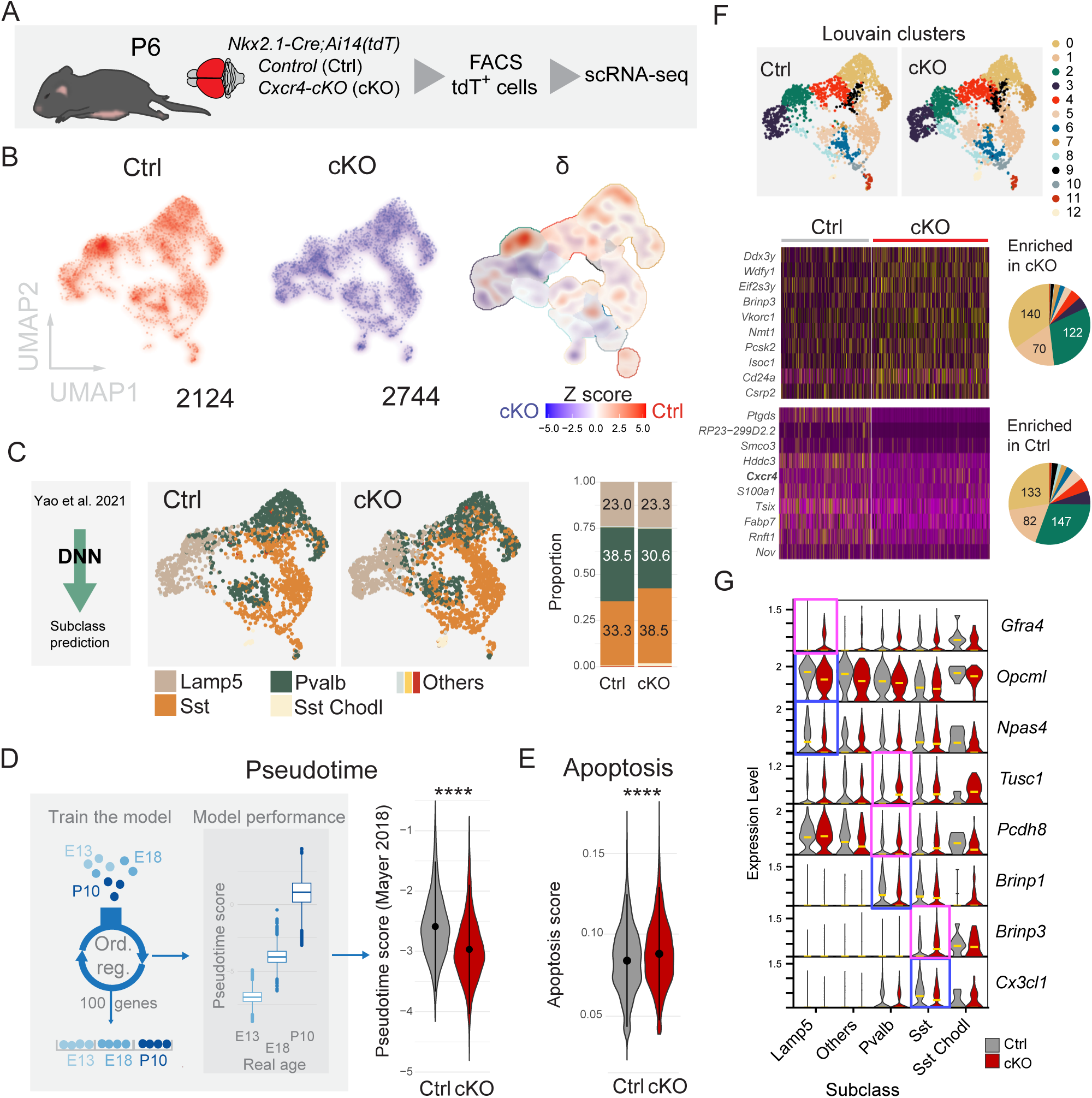
Positioning affects molecular identity and maturation of mCIN. **A**, Schematic illustration of the experimental strategy for single-cell RNA sequencing (scRNA-seq). **B**, UMAP visualizations shows single mCIN sequenced from control (Ctrl, red) and *Cxcr4*-cKO (cKO, blue) mice, with differences in cell density highlighted (δ). Numbers indicate total neurons sequenced per genotype. **C, Left**, UMAP representation of mCIN, color-coded by Deep Neural Network (DNN)-derived subclass labels. **Right**, Proportions of mCIN subclasses in control and *Cxcr4*-cKO samples. Note shifts in subclass composition. **D, Left,** Schematic illustration and cross-validation performance (boxplot) of the ordinal regression model used to predict the developmental pseudotime of mCIN, trained on the dataset provided in {Mayer, 2018 #63}. **Right**, Predicted pseudotime scores for control and *Cxcr4*-cKO mCIN. **E,** Apoptosis scores for control and *Cxcr4*-cKO mCIN, calculated using genes provided in {Park, 2020 #66}, and the UCell algorithm**. F, Top,** UMAP visualization of mCIN, color-coded by Louvain clusters. **Bottom**, Analysis of differentially expressed genes (DEGs), with selected up- and downregulated genes illustrated as heatmaps and the proportion of DEGs per cluster shown as pie charts. **G**, Violin plots show significantly up- (magenta box) or down-regulated (blue box) DEGs in *Cxcr4*-cKO cells across the indicated mCIN sublasses. **Statistics**: ****p <0.0001 as by Kolmogorov-Smirnov-Test (***D,E***).

We evaluated mCIN subclass composition by training a Deep Neural Network using the Allen Brain Atlas dataset (43, 44) as a reference to predict cell identity of our data (**Figures 3C** and **S5B**). While all major subclasses were present, there was a shift in the relative proportions of *Sst* and *Pvalb* mCIN subclasses, with a higher proportion of *Sst* relative to *Pvalb* in *Cxcr4*-cKOs, suggesting a partial change in molecular identity, probably driven by mispositioning. The *Lamp5* subclass remained unaffected. The shift in subclass ratio between *Sst* and *Pvalb*, but not *Lamp5 Lhx6*, was confirmed using MapMyCell as an alternative cell identity prediction method (**Figure S5C**).

To test if precocious invasion of the cortical layers influenced the developmental trajectory of mCIN in *Cxcr4*-cKOs, we trained an ordinal regression model using published developmental datasets (32, 45) to predict a relative ‘age’ score for our P6 mCIN. This revealed that *Cxcr4*-cKO mCIN exhibited a younger gene expression profile than control mCIN (**Figures 3D, S5D** and **S5E**), suggesting aberrant migration is connected to delayed mCIN maturation.

Given that the number of principal neurons determines the number of CIN in cortical circuits with CIN apoptosis as a main mechanism to adjust neuronal numbers (46, 47), we reasoned that aberrantly increased mCIN number in deep layers may promotes mCIN apoptosis in *Cxcr4*-cKOs. scRNA-seq indeed indicated a subtly elevated apoptotic pathway in *Cxcr4*-cKO mCIN, which most likely derived from *Sst* and *Pvalb* mCIN (**Figures 3E** and **S5F**). We then identified broader molecular alterations by performing differential gene expression analysis between control and *Cxcr4*-cKO mCIN within individual clusters and subclasses. This identified hundreds of DEGs (**Figures 3F**, **3G,** and **S5G; Table S2**) including subclass-specific DEGs previously associated with GABAergic neuron development and neurodevelopmental disorders, such as *Gfr4* (48), *Npas4* (49, 50), *Cx3cl1* (51), *Opcml* (52), *Pcdh8* (53), and *Brinp1* & *3* (54, 55).

Notably, as the mCIN transcriptome was virtually unaffected in E16.5 *Cxcr4*-cKOs, when the majority of mCIN are still under direct control of CXCR4 signaling, gene expression changes at P6 are developmental consequences rather than a direct effect of CXCR4 loss. Together, our RNA-seq results indicate that aberrant layering of mCIN during the perinatal period changes their molecular identity, developmental trajectory, and gene expression, highlighting a critical role of laminar positioning in CIN development.

### Altered network architecture in the neocortex and reduced mCIN numbers in the hippocampus at the mature stage

We next examined if mCIN subclasses were affected in *Cxcr4*-cKOs at P15, when the SST^+^ and PVALB^+^ mCIN can be distinguished by immunostaining. Like in the previous analyses, we chose the motor cortex as ROI. The mCIN population as a whole exhibited a slight downward shift (**Figure 4A**). This redistribution was driven by SST⁺ mCIN, which were reduced in superficial and increased in deep layers, and not by PVALB^+^ mCIN (**Figures 4B** and **4C**). The overall mCIN numbers were unchanged (**Figures 4A-4C**). We then investigated whether similar changes were present in other cortical regions. First, we chose the visual cortex for its caudal position as opposed to the rostral motor cortex. Assessments at P4, P15, and P30 (**Figures S6A – S6G**) revealed essentially similar layering defects as in the motor cortex (**Figures 4A-4C** and **S3A**). Notably, while the layering defect of the overall mCIN population partially recovered during the first postnatal weeks–with the mutant distribution converging toward control patterns **(Figures S6H** and **S6I),** the SST^+^ mCIN subpopulation was persistently shifted downward in the visual and motor cortex, suggesting this defect is permanent and present throughout the neocortex. Next, we turned to the hippocampus, as a medial limbic cortex area. At birth, this structure exhibited a reduced mCIN density in the stratum lacunosum moleculare (**Figure S7A)** – the hippocampal correlate of the neocortical MZ. This deficit transitioned into a persistent numerical reduction of SST^+^ mCIN across all hippocampal subregions by P30 (**Figures S7B** and **S7C**). Unlike in neocortex, SST^+^ mCIN did not show significant laminar redistribution in the hippocampus (**Figures S7D**). However, their reduced numbers led to a significant reduction in the innervation of the stratum lacunosum moleculare by SST^+^ axons **(Figures S7E–S7I)**.

**Figure 4.**
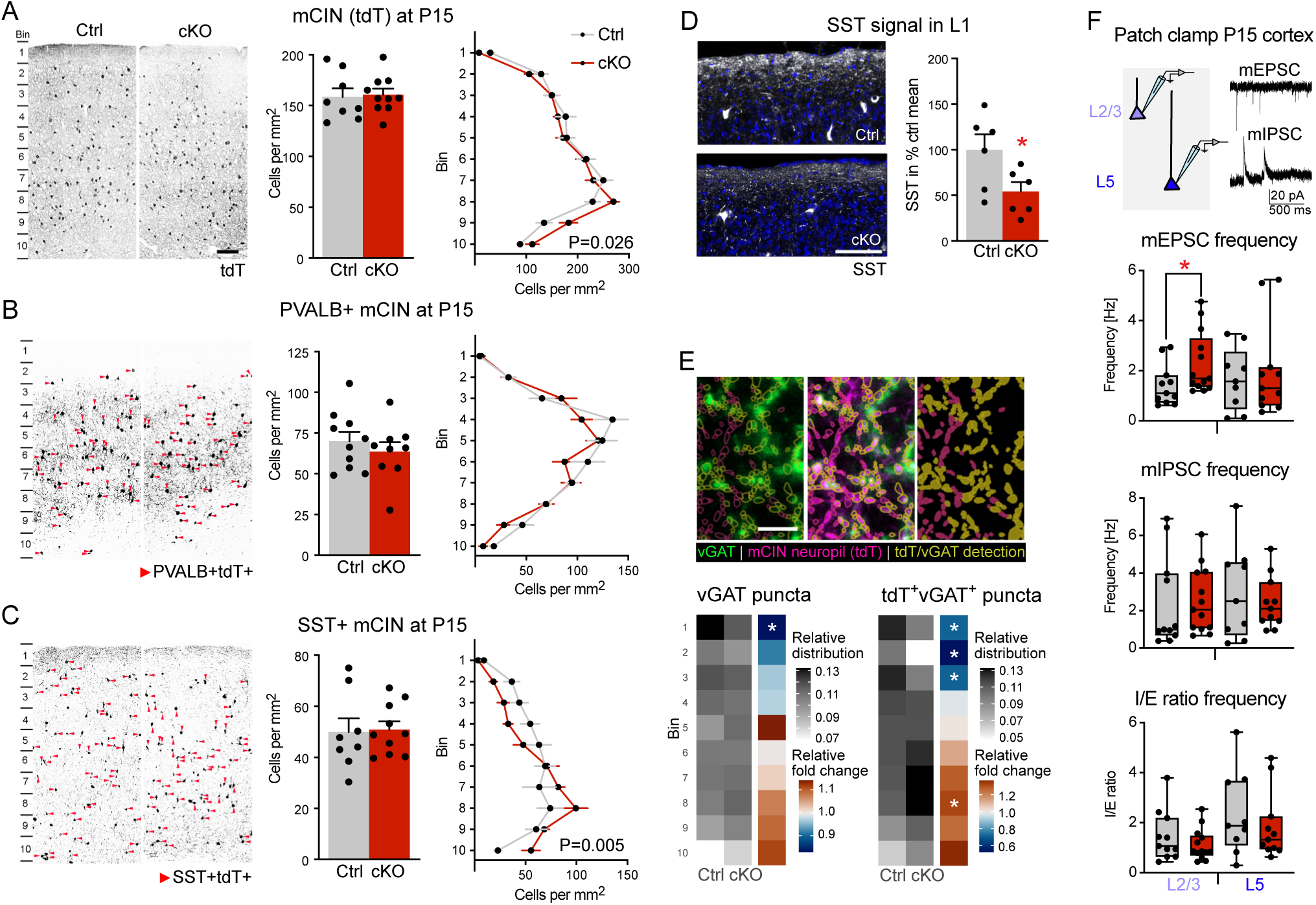
Shift of SST^+^ mCIN from superficial to deep cortical layers in *Cxcr4*-cKO mice. **A-C,** Confocal images demonstrate immunofluorescences for tdT (***A***), PVALB (***B***), and SST (***C***) in the motor cortex of P15 control and *Cxcr4*-cKO mice (*Nkx2.1*-Cre; *Rosa26*^LSL-tdT^). All PVALB^+^ and SST^+^ cells that are also tdT^+^ are highlighted by arrowheads. Bar and line graphs show the total number and the distribution of tdT^+^ (***A***), PVALB ^+^(***B***), and SST^+^ (***C***) mCIN across layers 1 to 6 of the motor cortex at P15. **D**, Immunofluorescence (left) and quantification (right) of SST in layer 1 (L1). **A-D,** Bar and line graphs show mean + SEM. Circles in bar graphs represent individual mice. **E, Top,** Detection of synaptic puncta using tdT (mCIN neuropil in magenta) and vGAT (in green) immunofluorescence (see Methods). **Bottom,** Heatmaps show the distribution of vGAT^+^ (left) and tdT^+^ vGAT^+^ (right) puncta across ten bins (layer 1 to 6) as relative distribution and as fold change (*Cxcr4*-cKO/ Ctrl). **F,** Miniature excitatory and miniature inhibitory postsynaptic potentials (mEPSC, mIPSC) recorded in pyramidal cells in layer 2/ 3 (L2/3) and layer 5 (L5) at P15 – P18. Box and whisker plots (minimum to maximum with median represented as horizontal line and individual cells as circles) show the frequency and the ratio of inhibitory and excitatory events for individual cells (I/E ratio frequency). **Statistics: A-C,** p-values for genotype-bin-interaction as by two-way ANOVA with repeated measures. Data depicted in the bar graphs are not significantly different as by Mann-Whitney test. **D,** *p = 0.0325 as by Mann-Whitney test. **E,** *p <0.1 as by beta regression model followed by simultaneous tests for general linear hypotheses for repeated measurements correction. **F,** *p = 0.0352, Mann-Whitney test. **Abbreviations:** cKO, *Cxcr4*-cKO; Ctrl, control; L1, cortical layer 1. **Scale bars:** 135 µm in ***A***, 100 µm in ***D***, 5 µm in ***E***. Data are available in **Table S5**.

The latter finding prompted us to assess SST^+^ terminals in the neocortex, where a large subpopulation of SST^+^ mCIN (Martinotti cells) sends their axons to layer 1 to provide inhibitory input to the distal dendrites of pyramidal cells (11). We noted that SST^+^ puncta were significantly reduced in layer 1 of *Cxcr4*-cKOs (**Figure 4D**). Since, unlike in the hippocampus, the overall number of SST^+^ mCIN was unchanged in the neocortex, this suggested altered axonal targeting. To test this assumption, we quantified the relative distribution of vGAT^+^ and tdT^+^ vGAT^+^ puncta across neocortical layers 1 to 6, showing a reduction in superficial layers including layer 1 and an increase in deep layers (**Figure 4E)**.

Given this finding, we assessed functional connectivity by recording miniature postsynaptic currents in pyramidal neurons in superficial and deep layers in *Cxcr4*-cKOs (**Figures 4F** and **S6J – S6L**). We chose the P15 – P18 period, because by that time the mature chloride equilibrium potential has been established (56), allowing us to record miniature excitatory and miniature inhibitory postsynaptic currents (mEPSCs and mIPSCs) from the same neuron. With the use of Tetrodotoxin, the recorded events resulted from spontaneous presynaptic neurotransmitter release and thus reflect the excitatory and inhibitory input to the recorded neuron. The frequency, amplitude, rise- and decay-times of mIPSCs were unchanged, whereas the frequency of mEPSCs was increased in superficial and unchanged in deep layers in *Cxcr4*-cKOs. Further parameters remained unaffected (**Figures 4F** and **S6J – S6L**). The overall increase in mEPSCs in superficial layers of *Cxcr4*-cKOs reflects an increase in excitatory synapses, perhaps as a result of altered network activity in the preceding postnatal period (57–59).

### Facilitated propagation of sensory-evoked activity in the mature cortex

We reasoned that the reduction of GABAergic terminals in combination with the increased number of excitatory synapses per pyramidal neuron present in superficial layers of *Cxcr4*-cKOs **(Figures 4E** and **4F)** might alter the propagation of neuronal activity between neocortical areas. To test this, we examined light-induced cFOS expression in various cortex areas as a readout for the propagation of stimulus-induced neuronal activity (60). Specifically, we kept P30 control and *Cxcr4*-cKOs for 18 h in complete darkness. Animals were then sacrificed either immediately (dark control group) or after a one-hour exposure to moderate light (light stimulus group). cFOS^+^ nuclei were counted in the primary visual, retrosplenial, and posterior parietal association cortex (i.e. the medial secondary visual cortex) (**Figures 5A**, **5C**, and **5D**). The retrosplenial and posterior parietal association cortices were included, because they receive inputs from primary visual cortex to integrate visual information for higher order cognitive functions (61). After darkness-adaptation, only few cFOS^+^ cells were present in the three areas in control and *Cxcr4*-cKOs. Light exposure increased the number of cFOS^+^ cells in the three areas, with no significant differences between genotypes in primary visual and retrosplenial cortex. The number of cFOS^+^ mCIN was not altered in the primary visual cortex of *Cxcr4*-cKOs following light-exposure (**Figure 5B**). Notably, in the parietal association cortex, light-induced cFOS expression was increased in *Cxcr4*-cKOs (**Figure 5D**), suggesting facilitated activity spread from primary sensory to association cortex. Collectively, these findings suggest that inhibitory containment of neuronal activity is affected in mature cortical circuits in *Cxcr4*-cKOs.

**Figure 5.**
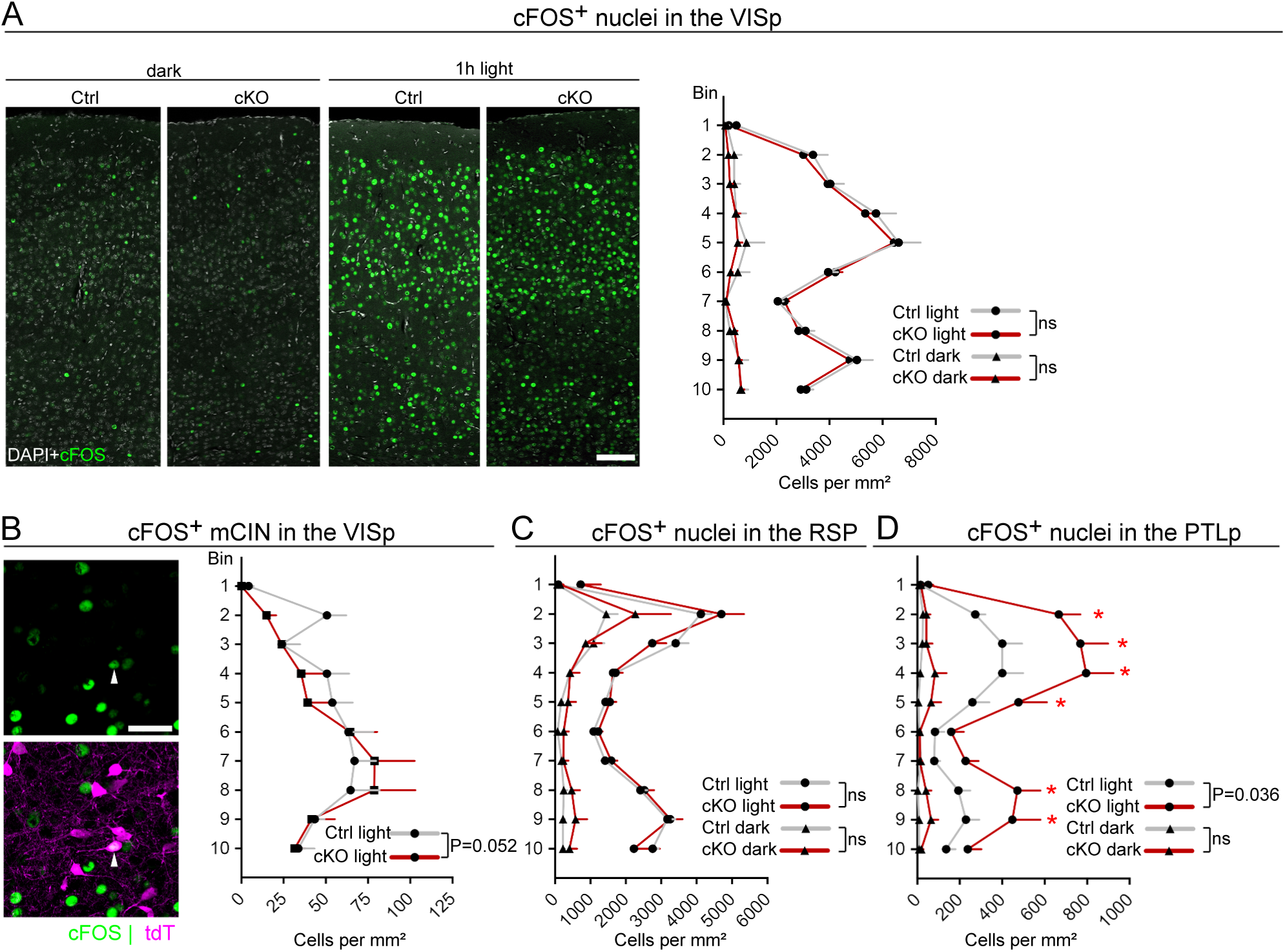
Light-induced cFOS expression is increased in the association cortex of *Cxcr4*-cKO mice. **A-D,** Images and graphs represent P30 control and *Cxcr4*-cKO mice (*Nkx2.1*-Cre; *Rosa26*^LSL-tdT^) after 18 h darkness (dark) or after 18 h darkness followed by 1 h moderate light (light). **A,** Confocal images demonstrate cFOS and DAPI in the primary visual cortex (VISp). The line graph shows the distribution of cFOS^+^ nuclei in the VISp. **B,** The high-magnification confocal image demonstrates a cFOS^+^ mCIN (arrowhead) in the VISp of a control mouse (light). The line graph shows the distribution of cFOS^+^ mCIN in the VISp. **C,D,** Line graphs show the distribution of cFOS^+^ nuclei in the retrosplenial cortex (RSP) (***C***) and posterior parietal association cortex (PTLp) (***D***). **Statistics:** p-values for genotype-bin-interaction as by two-way ANOVA with repeated measures; *false discovery rate (q) <0.1 as by two-stage linear step-up procedure (***A***-***D***). All graphs show mean + SEM. **Abbreviations:** cKO, *Cxcr4*-cKO; Ctrl, control; ns, not significant. **Scale bars:** 100 µm in ***A***, 40 µm in ***B***. Data are available in **Table S5**.

### Increased spatial spread of spontaneous activity in the visual cortex prior to eye opening

Spontaneous network activity in the early postnatal period is critical for establishing appropriate connectivity across developing cortical circuits and is constrained by GABAergic inhibition, which involves contributions from SST^+^ CIN limiting cellular recruitment during spontaneous network events (57–59). We therefore hypothesized that aberrant laminar positioning of mCIN in *Cxcr4*-cKOs impairs this inhibitory control, altering spontaneous network activity.

We thus performed transcranial wide-field Ca^2+^ imaging in the visual cortex of head-fixed mice prior to eye opening. We selected a period of physiological blindness (P3 to P5) and a period after the onset of retinal light-sensitivity (P10 to P12). We used pressure ejection to deliver the synthetic indicator OGB1 into the superficial layers of the primary visual cortex (**Figures 6A** and **6B**). For analysis, the ∼1 mm² field of view was divided into a grid of ROIs (**Figure 6C).** Spontaneous network activity occurred as discrete spatiotemporal clusters of Ca^2+^ transients (CaTs), alternating with silent periods (**Figures 6D** and **6E**). This discontinuous pattern was consistently observed in all mice (n=34) both in the early and late cohorts, in line with published data (62, 63). We detected Ca^2+^ clusters based on the fraction of active cells within a sliding window (Δt=0.13 s). Clusters exhibited robust developmental changes, including increased frequency and size, alongside decreased duration **(Figures 6F - 6L)**. Generalized linear modeling revealed a significant increase in cluster size in *Cxcr4*-cKOs across developmental stages (**Figures 6G - 6I**). In contrast, cluster frequency, amplitude, and duration (**Figures 6J - 6L**) were unaffected, indicating a selective increase of the spatial spread of spontaneous activity in *Cxcr4*-cKOs. To probe the timescale dependence of this effect, we systematically varied Δt from 0.13 to 2.16 seconds (**Figure 6E**: bottom). *Cxcr4*-cKOs showed increased spatial spread of Ca^2+^ clusters only at Δt=0.13–0.27 seconds (**Figure 6I**), indicating a specific enhancement of fast-timescale network recruitment, rather than slow-timescale coordination or overall activity levels.

**Figure 6.**
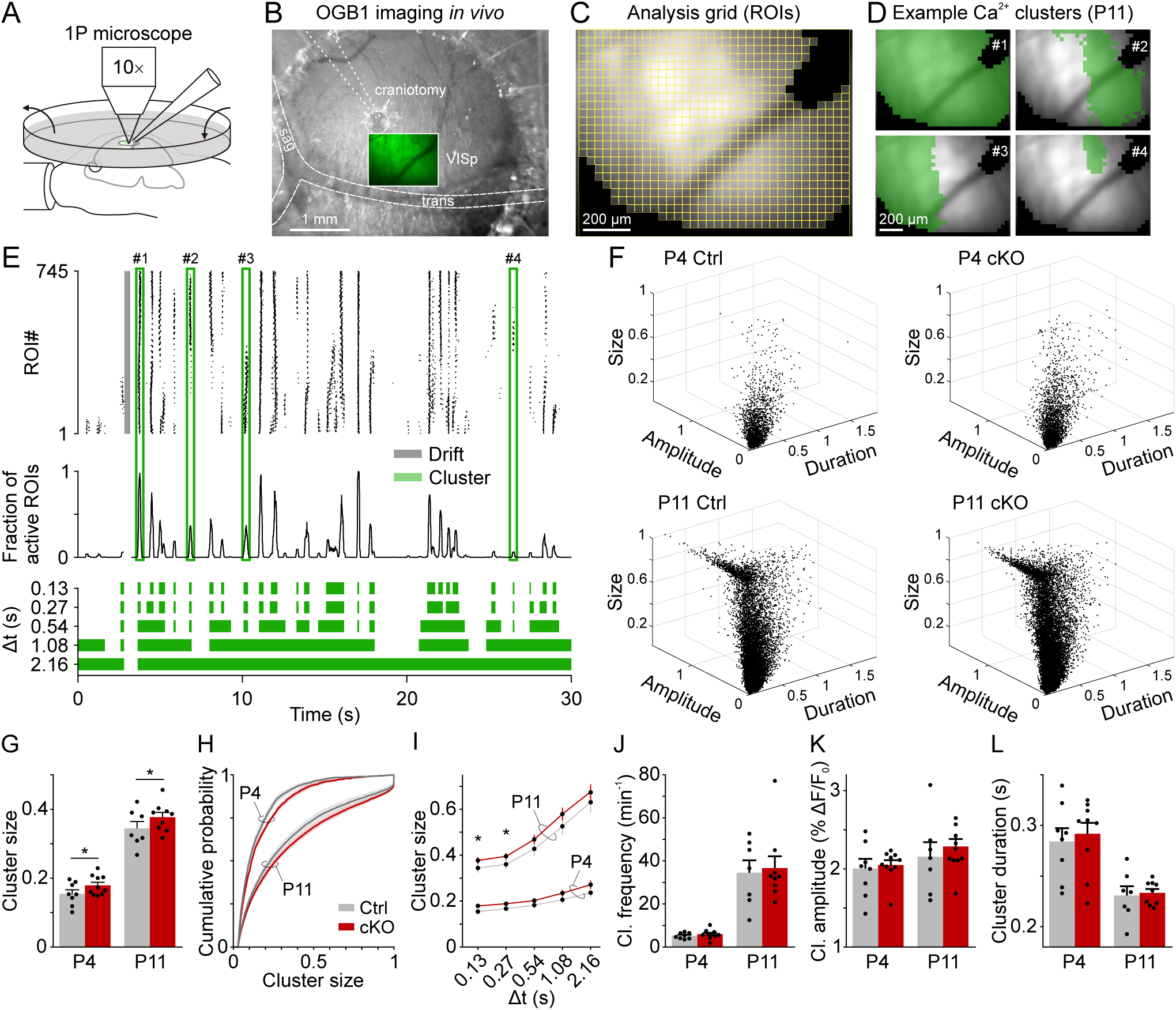
Spontaneous activity in the primary visual cortex (VISp) of *Cxcr4*-cKO mice before eye opening. **A,** Experimental setup for *in vivo* one-photon imaging under N_2_O anesthesia. **B,** Anatomical overview image showing the craniotomy site for OGB-1 dye application via pressure injection into the VISp. Green image represents the resting fluorescence signal. **C,** Resting fluorescence image of OGB-1, corresponding to the field of view shown in ***B***, overlaid with the grid of ROIs. ROIs marked in black were excluded from analysis due to the presence of large blood vessels or insufficient dye loading. **D,** Example Ca^2+^ clusters highlighted in panel ***E***. ROIs with detected Calcium transients are marked in green. **E, Top,** Raster plot of detected Calcium transients across ROIs over time. Grey-shaded areas indicate periods of animal movement (drift), which were excluded from the analysis. **Middle,** Fraction of active ROIs over time, calculated by summing Calcium transients within a moving window of 0.13 s. **Bottom,** Ca^2+^ cluster indicators computed for different temporal windows (Δt). **F,** 3D plots of all detected Ca^2+^ clusters in control and *Cxcr4*-cKO mice at P4 and P11, illustrating their size (fraction of field of view), amplitude (average % ΔF/F_0_) and duration (s). **G,** Cluster size is significantly increased in *Cxcr4*-cKO mice at both P4 and P11. **H,** Cumulative probability distributions indicate a shift toward larger cluster sizes in *Cxcr4*-cKO mice at both developmental stages. **I,** Average cluster size as a function of the temporal window Δt. **J-L,** No genotype-dependent differences were observed in Ca^2+^ cluster frequency (***J***), mean amplitude (***K***), or duration (***L***). Each circle represents an individual animal. Data are presented as mean ± SEM. *p <0.05. Data are available in **Table S5**.

During the first postnatal week, spontaneous activity in the primary visual cortex is dominated by low-spread clusters (L-events) reflecting retinogeniculate input, while high-spread, cortex-driven clusters (H-events) emerge shortly before eye opening (62, 64). To determine whether the observed increase in spatial spread preferentially affects one of these event types, we applied an iterative thresholding method to classify L- and H-events based on a three-dimensional feature space (cluster size, amplitude, duration) defined from control data at P3 – P4 (**Figures S8A** and **S8B**; see Methods). We found that L-events were significantly larger in *Cxcr4*-cKOs in both the early and late cohorts **(Figures S8C** and **S8D)**, while the size of H-events at P11—representing 15.4 ± 2.2% and 18.5 ± 1.4% of all clusters in control and *Cxcr4*-cKO mice—remained unchanged. H-events were rare at P3 – P4 and thus not analyzed at this stage. Our findings suggest that displacement of mCIN in *Cxcr4*-cKOs selectively disrupts the spatial confinement of retinally driven network bursts. To validate this conclusion, we applied a complementary bootstrapping and regression approach to clusters rank-ordered by size. Linear regression up to the kink point revealed a 23.8% and 13.2% increase in cluster size in *Cxcr4*-cKOs compared to controls at P3 – P4 and P10 – P11, respectively (**Figures S8E** and **S8F**).

In summary, aberrant laminar positioning of mCIN resulting from perturbed TM caused an increased horizontal spread of spontaneous activity in developing primary visual cortex. This effect was specific to fast-timescale network recruitment and selectively affected retinally driven L-events, consistent with spatial disinhibition due to impaired GABAergic signaling by displaced mCIN.

## Discussion

We established *Cxcr4*-cKO mice as a model of misrouted mCIN migration. Our results reveal that misrouting influences the layering and molecular identity of mCIN, as well as the synaptic connectivity and spontaneous activity of the cortical network during the early postnatal period. Further, while gross stratification defects of the mCIN population as a whole are gradually compensated, a permanent lack of SST^+^ mCIN persists in superficial layers and is associated with facilitated spread of sensory stimuli between neocortical regions. These findings connect CXCR4-guided mCIN migration to functional maturation of the cerebral cortex.

*Cxcr4-*cKOs exhibited normal long-term survival and breeding rates and no overt neurological or behavioral abnormalities during routine husbandry. This distinguishes our model from other CIN migration mutants presenting with developmental comorbidities or early lethality (27, 65, 66). These more severe phenotypes may be caused by motility defects resulting in severely reduced CIN numbers and/ or from disrupted CIN maturation. In our model, the direct *knockout*-effect is restricted mainly to layer-specific guidance during the tangential migration period. This is due to the transient expression profile of *Cxcr4* in mCIN (18, 35–37). Critically, although mCIN migrated via ectopic routes, they effectively colonized the neocortex of *Cxcr4*-cKOs by birth. Thus, *Cxcr4*-cKOs are a model to study functional consequences of early mCIN layering abnormalities, while numerical mCIN anomalies are restricted to distinct cortical areas (mainly limbic regions). Since the hippocampus is particularly remote from the mCIN origin (16), its reduced colonization indicates that combined speed-enhancing and guidance effects of CXCR4 support long-distance migration of mCIN as proposed (14, 20, 38–40).

Our findings in *Cxcr4*-cKO mice show that proper guidance during the embryonic period is crucial for mCIN to populate the neonatal MZ, and that especially late-born mCIN colonize the late-forming superficial cortical layers from this niche during the first postnatal week. Given that CIN linger in the MZ for days (14), we propose that the MZ serves an intermediate station for mCIN until late-arriving principal neurons establish the superficial layers. The failure to populate the MZ reservoir leads to a deficit of mCIN in the superficial layers of *Cxcr4-*cKOs. Our BrdU-labeling study indicates that the radial migration, which sorts mCIN into the appropriate layer according to their birth-date (42), is largely CXCR4-independent. Nevertheless, it corrected the early misplacement of mCIN in *Cxcr4*-cKOs only partially. In particular, a significant proportion of late-born mCIN remained misplaced in deep layers by P8, consistent with mCIN transplantation studies (13, 33). Our scRNA-Seq analysis at P6 revealed a more immature transcriptional profile of *Cxcr4*-deficient mCIN, suggesting that they remained in a prolonged radial migration-mode while attempting to reach their target layers.

The layering abnormalities of mCIN in *Cxcr4*-cKOs decreased further between P8 and P15, indicating the presence of compensatory mechanisms following the conclusion of active mCIN migration. mCIN apoptosis might be one such mechanism as suggested by scRNA-seq. Since apoptotic elimination of mCIN adjusts their numbers to those of principal neurons (46, 47, 67), apoptosis may serve as a compensatory mechanism in *Cxcr4*-cKOs particularly in layers with an excess of mCIN.

Since the identity of mCIN is not fully established by the end of the migration period (32), it is plausible that abnormal environmental signals due to misplacement may affect mCIN maturation. A series of subclass-specific DEGs associated with GABAergic neuron development and neurodevelopmental disorders detected in mCIN from P6 *Cxcr4*-cKOs supports the notion that positioning instructs mCIN maturation. We further identified a higher proportion of *Sst*-relative to *Pvalb*-class-mCIN in P6 *Cxcr4*-cKOs. However, immunohistochemistry showed a normal density of PVALB^+^ mCIN and a normal SST/ PVALB ratio at P15. We assume that the *Sst*/ *Pvalb* imbalance at P6 reflects a delay in the maturation of PVALB^+^ mCIN. Mechanistically, this could be caused by abnormal positioning and protracted migration behavior. In addition, altered network activity could delay PVALB maturation in *Cxcr4*-cKOs, because activity of SST^+^ mCIN can delay PVALB maturation (68).

At the mature stage, the most prominent anatomical defect in *Cxcr4*-cKOs was a shift of SST^+^ mCIN from superficial to deep neocortical layers, accompanied by a reduction of SST^+^ axonal puncta in layer 1. This likely reflects the dependence of a subset of SST^+^ mCIN (Martinotti cells) on MZ migration for axonal projection into neocortical layer 1 (69). Given these defects, the reduction in tdT^+^ vGAT^+^ puncta (representing GABAergic terminals), and the increase in excitatory synapses per pyramidal neuron in superficial layers of *Cxcr4*-cKOs, we hypothesized that communication between neocortical areas might be changed in *Cxcr4*-cKOs. In fact, propagation of light-induced neuronal activity was facilitated between primary visual and posterior parietal association cortex, while no such facilitation was present in retrosplenial cortex. This differential response may stem from stronger projections from the primary visual to the posterior parietal association cortex compared to retrosplenial cortex (61). The facilitated stimulus propagation in *Cxcr4*-cKOs likely is a consequence of altered cortical network architecture including a lack of inhibition in superficial layers.

A major finding of this study is that aberrant laminar positioning of mCIN alters *in vivo* network dynamics in the visual cortex prior to eye opening, manifested as an increased lateral spread of L-events. As spontaneous activity in the early postnatal period is critical for establishing appropriate connectivity across developing cortical circuits (57–59), this finding links guided CIN migration to cortical network maturation. Spontaneous local network events are triggered by retinogeniculate input (64) and are electrophysiologically characterized by spindle bursts, which interact with retina-independent H-events to jointly shape network connectivity via activity-dependent plasticity rules (64, 70). Depolarizing GABA normally imposes inhibitory constraints on developing visual networks (57, 71). Our data therefore indicate that TM deficits lead to functional disinhibition, consistent with reduced axonal SST signals in layer 1 and decreased mCIN-derived synapse densities in superficial layers. Notably, mIPSCs in layer 2/3 and layer 5 pyramidal neurons shortly after eye opening were unaffected, suggesting that the inhibitory deficit more likely reflects a spatial redistribution of GABAergic input rather than a reduction in overall inhibitory strength. Glutamatergic signaling may also contribute, as mEPSC frequency in layer 2/3 pyramidal neurons was slightly increased in *Cxcr4*-cKO mice. Our findings indicate that mCIN constrain the spatial spread of L-events, whereas H-events remain unaffected. This is consistent with previous work showing that chemogenetic silencing of SST interneurons selectively increases cell participation in L-events, allowing them to propagate farther across the cortex (59).

Misrouting of tangentially migrating CIN results in significant anatomical, molecular, and functional disturbances during the early postnatal period when the cortical network is established and refined by spontaneous network activity. Although gross anatomical defects in mCIN layering are compensated postnatally, cortical function is permanently affected, as evidenced by facilitated propagation of sensory-evoked activity at the mature stage. By linking perturbed guidance of migrating CIN to abnormal spontaneous network activity in the early postnatal period, we provide mechanistic insights into how developmental defects may contribute to altered cortical function in neurodevelopmental and neuropsychiatric disorders.

## Materials and Methods

### Mice

Animal husbandry and surgery were in accordance with institutional and EU or national guidelines for animal use, approved by the competent authority (license numbers UKJ-22-020 an UKJ-20-014) and supervised by the institutional veterinarians. Mice were kept on C57BL/6j background under temperature-controlled conditions (20-24°C) with a 12 h dark-light cycle and free access to food and water. Developmental stages of embryos are stated in the results section and figure legends. Noon of the day after mating was considered E0.5. Animals were used irrespective of sex and allocated to experimental groups only according to their genotype. The following mouse lines were used: *Cxcr4*-LoxP, Jackson, stock no. #008767; *Nkx2.1*-Cre, Jackson, stock no. 008661; Ai14 (*Rosa26*^LSL-tdT^), Jackson, Stock no. #007914. For histology and RNA-sequencing, *Nkx2.1*-Cre; *Cxcr4*^LoxP/LoxP^; *Rosa26*^LSL-tdT/wt^ mice were used as *Cxcr4*-cKOs and *Nkx2.1*-Cre; *Cxcr4*^LoxP/wt^; *Rosa26*^LSL-tdT/wt^ or *Nkx2.1*-Cre; *Cxcr4*^wt/wt^; *Rosa26*^LSL-tdT/wt^ mice were used as controls.

### BrdU pulse-labeling

*Cxcr4*-cKO and control mice (*Cxcr4*^LoxP/wt^) were generated as littermates using a *Cxcr4*^LoxP/LoxP^ × *Cxcr4*^LoxP/wt^ breeding scheme. Pregnant females received BrdU i.p. (100 mg/kg) at 8:00 and 16:00 either on E12 or on E15. Offspring were collected on P0 and P8. For a given litter, a similar number of *Cxcr4*-cKO and littermate controls was included into the experimental cohort.

### Histochemistry

*Lhx6 in situ* hybridization was performed as described (25). For immunohistochemistry and detection of BrdU, whole embryos (E13.5), heads (E16.5), and postnatal brains were fixed overnight in 4% PFA, PBS, pH7.4 and processed as described (72). Antibodies, fluorescent Streptavidins, and dilutions are detailed in **Table S3**. For detection of BrdU, sections were treated for 10 min in 1N HCl on ice, for 10 min in 2N HCl at RT, and for 20 min in 2 N HCl at 37°C. Section were then transferred for 12 min to 0.1M borate buffer (pH 8.5). After 15 min treatment with TPBS-T, a blocking step was performed for 1h in TPBS-T containing 3% BSA and 1 mol/l Glycin. Application of anti-BrdU and anti-RFP primary antibodies and detection were performed as described above (72). Representative micrographs demonstrating *Lhx6* and immunohistochemistry were captured using Axio.Imager A1 (Zeiss) connected to Gryphax camera (Jenoptik) and LSM900 (Zeiss), respectively. Images were processed with Adobe Photoshop (v6). Micrographs of control and *Cxcr4*-cKO samples of the same experiment were always processed in the same way.

### Image Analysis

#### Density and layering of mCIN and BrdU^+^ cells

Micrographs were taken with LSM510 Meta (Zeiss), LSM900 or Axio.Imager A1 as detailed in **Table S4**. Specimens of the same experiment were captured with identical imaging conditions. Using Fiji software, images scaled in µm were rotated so that the pia mater (neocortex) or the ependyma of the lateral ventricle (hippocampus) was oriented horizontally at the top. After rectangular cropping, the upper and lower borders of the ROI were labeled using the Fiji multi-point tool. Marker coordinates were exported and interpolated to define the y-coordinates of the upper and lower border of the ROI in 1 µm steps. Cells were labeled using the Fiji cell multi-point tool, switching between channels when multiple markers were considered (e.g. tdT, DAPI, and SST). When counting BrdU^+^ and BrdU^+^/ tdT^+^ cells in animals receiving BrdU on E12.5, only strongly labeled cells were considered to focus on cells undergoing the final division. Coordinates of the labeled cells were exported. For bin assignment of a cell, the y-coordinate of a cell was expressed relative to the y coordinates of the upper and lower border at the celĺs x coordinate and and defined as its relative depth (RD; upper border = 0% RD, lower border = 100 % RD). Depending on the experimental question, four to ten equally sized bins were defined. Cells were assigned to bins according to their RD. The absolute cell density (in cells per mm^2^) of an individual bin was calculated as follows: 10^6^ × number of bins × number of cells per bin / area of the ROI (the area in µm^2^). To determine the relative cell density of a single bin, the number of cells per bin was divided by the total number of counted cells. To establish relative distribution curves across bins, kernel density estimates (KDE) were calculated from the RD of all cells in an experimental group (R package ggplot2 v.3.5.1). All assessments were performed in frontal sections. Layering of tdT^+^ cells in the E16.5 neocortex was examined at the level of the anterior commissure in the somatosensory cortex. ROIs at postnatal stages are detailed in the Results section.

#### Density and layering of cFOS^+^ cells

Images were taken with Axio.Imager A1. ROIs were defined as described in the previous paragraph. cFOS^+^ cells were counted by the Fiji-plugin StarDist with standard settings. For bin-assignment, the x- and y-coordinates of the counted cells were exported and processed as described above. tdT^+^ cFOS^+^ cells were counted manually and assigned to bins as described above.

*Layering of VGAT^+^ and tdT+/vGAT+ puncta* (representing GABAergic synapses): Images encompassing layers 1 to 6 of the VIS were acquired using LSM900 and a 63x/1.4 oil objective (Plan-Apochromat, Zeiss). vGAT or tdT immune-positive synaptic sized objects (puncta) were detected across the entire cortical column using the StarDist ImageJ plugin or StarDist extension in Qupath (73) with parameters *normalizePercentiles (1, 99), .threshold(0.5), .pixelSize(0.02)*. To specifically identify GABAergic synapses derived from the mCIN population, an object classifier was trained to distinguish between vGAT^+^tdT^+^(representing GABAergic synapses, yellow in **Figure 4E** and vGATtdT^+^ (representing other mCIN neuropil components, magenta in **Figure 4E**). Puncta were assigned to bins according to their RD. To establish the relative distribution, the puncta count per bin was divided by the sum of all puncta. Further, the fold change was calculated for *Cxcr4*-cKO vs. control (*Cxcr4*-cKO/Ctrl ratio).

#### Statistical analyses of absolute cell densities

Calculations were performed using GraphPad Prism 10 for macOS (v10.5.0) using Mann Whitney test for a single ROI and 2way ANOVA followed by two-stage linear step-up procedure of Benjamini, Krieger, and Yekutieli for multiple ROIs or bins. p values <0.05 (Mann Whitney test, 2way ANOVA) and false discovery rates (q values) <0.1 for genotype or genotype x bin interaction were considered as statistically significant.

#### Statistical analyses of relative distributions

Distributions of cells and synapses were statistically compared between genotypes using R Statistical Software (v.4.4.2, R Core Team 2024) and a beta regression model (betareg package v.3.1.4). To account for repeated measurements, the model was followed by Simultaneous Tests for General Linear Hypotheses (glht function, multcomp package v.1.4.25; reference (74)).

### Bulk RNA sequencing

#### Fluorescence-activated sorting of mCIN from E16.5 cortices

Dissected cortices from E16.5 *Cxcr4*-cKOs and littermate controls were dissociated with the neural tissue dissociation kit P and gentleMACS dissociator (Miltenyi). Suspensions were passed through a 100 μm cell strainer. Cells were collected in FACS buffer, washed and resuspended in FACS buffer containing 5% BSA. Samples were immunostained with anti-CD140a-APC antibody (1:100 in FACS buffer) for 30 min at 4°C and passed through a 40 μm strainer immediately before sorting and analyzing with a BD FACSAria Fusion and BD FACSDiva (v8.0.2) and FlowJo (v10.1, LLC) software. After side scatter (SSC-A) and forward scatter (FSC-A) gating, gating on single cells and dead cell exclusion (Zombie Violet), a gate was set on CD140^-^ (PDGFRA^-^) tdT^+^ cells to sort mCIN into RLT buffer (QIAGEN) to ensure immediate stabilization of RNA.

#### Library construction and sequencing

Total RNA was extracted according to the manufacturer’s instructions (QIAGEN). mRNA was isolated, primed using poly-T oligonucleotides, and then converted into cDNA via SMART reverse transcription. Pre-amplification was performed using SMART ISPCR, followed by fragmentation with the Nextera XT DNA Library Preparation kit (Illumina), amplification, and indexing. The quality and fragment size distribution of the cDNA libraries were assessed using a high-sensitivity D5000 assay on the Tapestation 4200 system (Agilent Technologies), and concentrations were determined with a Qubit high-sensitivity dsDNA assay (Thermo Fisher Scientific). Libraries passing QC were pooled equimolarly and subjected to sequencing. Sequencing was performed on an Illumina NovaSeq platform in a paired-end configuration with 50 bp read length. The raw transcriptome files and count data will be deposited in NCBI’s Gene Expression Omnibus (GEO) and will be accessible under GEO Series accession number x. Transcript abundances were quantified using *kallisto* (75) against the Gencode M16 mouse transcriptome annotation (https://www.gencodegenes.org/mouse/release_M16.html). The resulting transcript-level estimates were summarized to the gene level and normalized with DESeq2 (v1.46.0) (76) using default size factor estimation, correcting for sequencing depth across samples. Lowly expressed genes with fewer than 10 total counts were excluded from further analysis. Differential gene expression was determined with the Wald test implemented in DESeq2, and p-values were adjusted for multiple testing using the Benjamini–Hochberg method. Genes with an adjusted p-value ≤0.1 were highlighted as significantly differentially expressed.

### scRNA-seq

#### Sample collection, RNA-seq library preparation and sequencing

Brains of six P6 controls and four *Cxcr4*-cKOs (both sexes) were collected in ice-cold Hibernate-A Medium (ThermoFisher, A1247501) with 10% FBS and B-27 supplement (ThermoFisher, 17504044). Cortices were manually dissected. A papain dissociation system (Worthington, LK003150) was used as described (77) on the gentleMACS Octo Dissociator (Miltenyi Biotec) to generate a cell suspension. tdT^+^ cells were sorted on BD FACSAria III (BD FACSDiva Software, v.8.0.2) with a 100 μm nozzle and subjected to downstream processing on the 10x Genomics Chromium platform. After sorting in PBS (Lonza, 17-516) with 0.02% BSA (B9000, NEB), ∼12’000 individual cells per sample were loaded onto a 10X Genomics Chromium platform for gel beads-in-emulsion and complementary DNA generation, carrying cell- and transcript-specific barcodes using the Chromium Single Cell 3’ Reagent Kit v.3.1 with Feature Barcoding technology (10X Genomics, PN-1000121) following the manufacturer’s protocol (document no. CG000205, 10X Genomics). 10x Genomics libraries were sequenced on an Illumina NovaSeq at the Genomics Core Facility of the Helmholtz Center Munich. Sequencing reads in FASTQ files were aligned to a reference transcriptome (mm10-2.1.0) and collapsed into UMI counts using the 10x Genomics Cell Ranger software (version 3.0.2).

#### Single cell analysis

All bioinformatics were performed using R Statistical Software (v.4.4.2, R Core Team 2024) and Bioconductor packages as described below. Graphs and heatmaps were plotted with the packages ggplot2 (v.3.5.1) and Seuratv5 (78) plotting functions.

#### Quality control

Libraries were preprocessed using a minimum of 1’500 genes detected, below 20% mitochondrial gene ratio cut-off and DoubletFinder (79) for doublet removal. Subsequently, counts were normalized and corrected for sequencing depth using Seurat’s *NormalizeData* or *SCTransform* function. Off-target cells were excluded by assigning cellular identity using a DNN preditions (44) (see below). Interneurons were analyzed using Seurat functions *SCTransform*, *RunPCA*, *RunUMAP*, *RunTSNE*, *FindNeighbors*, *FindClusters* (resolution=1) with default parameters and dims=1:30.

#### Pseudotime

Cells were pseudotime-aligned based on indicated reference and the R package bmrm (version 4.4) (80). Briefly, a regularized ordinal regression method (**Fehler! Verweisquelle konnte nicht gefunden werden.**) was used to predict cellular age within the present data set by training on published data from reference (32) (E13.5, E18.5, and P10, GSE104156) and reference (45) (P2, P10, P28, GSE165233), respectively. The linear weight of the model is used to rank genes according to their ability to predict each cell in time, followed by selection of 100 core genes (top 50 and bottom 50) for each model. The pseudotime score for each cell in the present dataset was then calculated by multiplying the log2-normalized gene counts by the trained core gene weights derived from each ordinal regression model.

#### Cell identity prediction

The Deep neuronal network (DNN) used to predict cell types was implemented in R with torch (0.12.0), luz (v0.4.0) and scml (v1.8.0) packages. The model accepted log-normalized gene counts as input and outputted prediction scores for N cell types. Each cell was assigned the type with the highest score. The sequential architecture included three layers. First, an input layer with dropout (p=0.5) was connected to a dense layer of 4,096 nodes, which calculated the cosine similarity with the node weights. This was followed by a hidden layer using dropout (p=0.25), a linear layer with 256 nodes, and a ReLU activation. The final output layer used dropout (p=0.25) and a linear layer with N nodes. The model was trained in two phases with a Multi-Margin criterion and the Adam optimizer, with a geometrically decreasing learning rate. The first layer was pruned after the initial phase, retaining the 100 most predictive inputs per node (50 highest and 50 lowest weights). Cortical cell types were used to train both a subclass and a subtype model.

Cell types were also predicted using the MapMyCells tool (RRID:SCR_024440), part of the Allen Brain Cell Atlas. The prediction was performed according to the instructions provided on the tool’s homepage (https://knowledge.brain-map.org/mapmycells/process)

*Apoptosis scoring:* Apotptic score were calculated using UCell package (81) and genes derived from reference (82) (30 genes), GO:0043065 (positive regulation of apoptotic process, 1016 genes) and GO:2001244 (positive regulation of intrinsic apoptotic signaling pathway, 87 genes).

#### DEGs

DEGs were identified by the Seurat function *FindAllMarkers* and *test.use = “MAST”* with default parameters (MAST v.1.24.1)(83). The top genes were plotted using *DoHeatmap* function.

### Patch-clamp recordings in acute brain slices

#### Preparation of acute cortical brain slices

*Cxcr4*-cKOs and heterozygous control littermates were decapitated during isoflurane anesthesia. The brain was removed in ice cold cutting ACSF (40 mM NaCl, 25 mM NaHCO3, 10 mM glucose, 150 mM sucrose, 4 mM KCl, 1.25 mM NH2PO4, 0.5 mM CaCl2, 7 mM Mg2Cl; purged with 95% O2 / 5% CO2, pH 7.35). The olfactory bulb was cut in the coronary plane and the hemisphere was glued with the cut face onto the probe-holder with superglue (UHU). 300 µm thick coronal slices were made with a vibratome (Leica VT1200S) with an amplitude of 1 mm and a velocity of 0.5 mm/s. Slices were transferred into an incubation beaker with cutting ACSF at 34°C for 1 hour and then transferred into another incubation beaker with recording ACSF (125 mM NaCl, 25 mM NaHCO3, 25 mM glucose, 2.5 mM KCl, 1.25 mM NaH2PO4, 1 mM MgCl2,2mM CaCl2, purged with 95% O2 / 5% CO2) for at least 1 hour before recordings at RT.

#### Patch-clamp recordings

Brain slices were transferred into a recording chamber, perfused with recording ACSF with 1 µM TTX (Tocris). Whole-cell recordings from cortical pyramidal cells were obtained with recording electrodes pulled from thick-walled borosilicate glass (2.0 mm o.d.) and filled with intracellular recording solution (135 mM CsMeSO4, 3 mM CsCl, 10 mM HEPES, 0.2 mM EGTA, 0.1 mM spermine, 2 mM QX-314-Br, 2 mM Mg2-ATP, 0.3 mM Na2-GTP, 10 mM phosphocreatine, pH 7.25, osmolarity 290 mOsmol, liquid-junction potential of 9 mV was corrected during recordings). mEPSCs and mIPSCs of visual cortex pyramidal cells were recorded in L2/3 (100 – 300 µm from pia mater) and L5 (100 – 300 µm from corpus callosum/ cortex boundary). Recordings were conducted for 100 s at holding potentials of −71 mV (reversal potential for GABAA receptors) for mEPSCs and +5 mV (reversal potential for AMPA and NMDA receptors) for mIPSCs. Series and input resistance was monitored during the recording time. Recordings were discarded if series resistance was higher than 25 MOhm or changed by more than 20% during the recordings. Series resistance was calculated from the peak current of a 5 mV voltage step in voltage-clamp. Input resistance was calculated from the steady state current of the same 5 mV voltage step. mEPSCs and mIPSCs were analyzed by MiniAnalysis Software (Synaptosoft) using a 1 kHz low-pass Butterworth filter at a detection threshold of 10 pA.

#### Surgical preparation, anesthesia and animal monitoring for *in vivo* imaging

Thirty minutes prior to surgery, animals received a subcutaneous injection of metamizole (Novacen, 200 mg/kg) for systemic analgesia. Animals were then placed onto a warm platform and anesthetized with isoflurane (3.5% for induction, 1–2% for maintenance) in pure oxygen (flow rate: 1 l/min). The skin overlying the skull was disinfected and locally infiltrated with 2% lidocaine (s.c.) for local analgesia. Scalp and periosteum were removed, and a custom-made plastic chamber with a central borehole (Ø 3–4 mm) was fixed on the skull using cyanoacrylate glue (UHU). The recording chamber was secured to the microscope stage and continuously perfused with oxygenated artificial cerebrospinal fluid (ACSF) containing (in mM): 125 NaCl, 4 KCl, 25 NaHCO_3_, 1.25 NaH_2_PO_4_, 2 CaCl_2_, 1 MgCl_2_ and 10 glucose (pH = 7.4 at 35–36 °C; ACSF flow rate ∼3 ml/min).

For dye loading, a craniotomy was performed above the left occipital cortex using a 27G needle **(Figure 6b)**, with careful preservation of the dura mater. Multicellular bolus loading of the Ca²⁺-sensitive indicator Oregon Green 488 BAPTA-1 AM (OGB-1, 500 μM) was performed by pressure injection through a glass micropipette (resistance: 4–5 MΩ) at a depth of 350–450 μm. Injections were performed over 10 minutes at a pressure of 8–9 PSI. Imaging commenced one hour post-injection to allow for intracellular de-esterification of the indicator dye.

During *in vivo* recordings, body temperature was continuously monitored and maintained at close to physiological values (36–37°C) by means of a heating pad and a temperature sensor placed below the animal. Respiratory activity was continuously monitored using a differential pressure amplifier (Spirometer Pod and PowerLab 4/35, ADInstruments). Upon completion of the surgical procedures, isoflurane was withdrawn and gradually replaced by nitrous oxide (N₂O) to achieve a final N₂O/O₂ ratio of 3:1 (flow rate: 1 l/min). Data acquisition began 60 minutes after complete withdrawal of isoflurane. At the end of each experiment, animals were euthanized by decapitation under deep isoflurane anesthesia (5%).

#### Wide-field epifluorescence microscopy

One-photon excitation of the Ca^2+^ indicator OGB1 was achieved using a xenon arc lamp (Lambda LS, Sutter Instrument), coupled via a liquid light guide to the epifluorescence port of a Movable Objective Microscope (Sutter Instrument). Excitation light was band-pass filtered at 472/30 nm, and intensity was adjusted to maintain baseline fluorescence within the mid-range of the camera’s dynamic range. Emitted fluorescence was separated at 495 nm and long-pass filtered at 496 nm (AHF Analysentechnik). Fluorescence images were acquired using a 10×/0.3 NA water immersion objective (Zeiss) and a 12-bit Rolera-XR camera (QImaging) controlled via Winfluor 3.7.5 (Dr. John Dempster, University of Strathclyde, Glasgow). Using 4×4 hardware binning (174×130 pixels), the sampling rate was ∼74.3 Hz. For V1 recordings the field of view was 1,160×867 µm. Imaging was performed transcranially through the intact skull in both age groups for approximately 45 minutes per session. The focal plane was set to 200–300 µm beneath the skull surface, corresponding to cortical layer 2/3 in neonatal mice.

#### Analysis of Ca^2+^ imaging data

Ca^2+^-imaging sequences were processed using custom MATLAB scripts. To improve signal to noise ratio raw images were spatially binned (5×5 pixels), yielding a grid of 34×26 = 884 ROIs (ROI dimensions: 33.35×33.35 µm). ROIs overlaying large blood vessels or exhibiting poor OGB1 labeling were excluded from further analysis.

#### Motion correction

Residual motion artifacts (“drift”) were identified semi-automatically by exploiting vascular landmarks. Specifically, each frame was filtered using the Frangi vesselness filter (FrangiFilter2D, D. Kroon, University of Twente) to accentuate high-contrast vascular structures. We then computed the Pearson correlation between each Frangi-filtered frame and a template image defined as the pixel-wise median of the first 2,200 filtered frames. The resulting correlation trace was inverted (mirrored) to detect local maxima, corresponding to potential motion events, using UFARSA (a general-purpose peak-detection algorithm, (84)). For each candidate peak, a median-ΔF/F₀ image was computed over five frames centered around the peak. When drift candidates were visually confirmed to be movement-related, 44 frames (∼0.6 s) centered on each motion-induced peak were discarded for subsequent analysis.

#### Detection of Ca^2+^ transients (CaTs)

After drift removal, fluorescence-versus-time traces (F(t)) were extracted for each ROI. Baseline fluorescence (F₀) was computed as the median of all F(t) values; ΔF/F₀ was then calculated as (F – F₀)/F₀. CaTs were detected on each ROI’s ΔF/F₀ trace using UFARSA (noiseSTD = 3.5), yielding a binary activity vector for each ROI (1 = CaT detected, 0 = no CaT).

#### Ca^2+^ clusters

Ca^2+^ clusters were operationally defined as follows: (1) CaTs in individual ROIs were classified as cluster-related if they fell into a time interval Δt during which the fraction of active ROIs (i.e., ROIs with >= 1 detected CaT) was >= 3%. (2) Neighboring cluster-related CaTs were assigned to the same cluster if both shared a time interval Δt during which the fraction of active cells was >= 3%, otherwise to separate clusters. Cluster duration was computed as the time difference between the last and the first CaT belonging to a given cluster. Cluster size was defined as the fraction of active ROIs for a given Ca^2+^ cluster. The Ca^2+^ cluster amplitude was calculated as the average amplitude of each Ca^2+^ transient defined as the maximum difference in the ΔF/F_0_ trace within the Ca^2+^ cluster duration.

#### Classification of L- and H-events

To distinguish low-synchronicity (L-events) versus high-synchronicity (H-events) network events (59), we employed a percentile-based iterative thresholding method in the three-dimensional feature space (cluster size, amplitude, duration).

The procedure was as follows:

1. Reference Distribution: P4 control animals defined the reference dataset. From their Ca²⁺ clusters, we derived robust estimates of median and median absolute deviation (MAD) for each of the three features.
2. Iterative Thresholding: Using a 98th-percentile criterion, we iteratively refined a multivariate threshold hyper-surface based on median/MAD until convergence.
3. Projection and Classification: Each Ca²⁺ cluster from every experimental group was projected into this reference feature space. Clusters falling below the threshold hyper-surface were labeled as L-events, while those exceeding it were labeled as H-events.
4. Subgroup Analysis: Finally, cluster-specific metrics were computed separately for L-and H-events, permitting direct comparison of *Cxcr4*-cKO versus control mice within each event category and across developmental stages. *Scaling of Cluster Size in Cxcr4-cKO.* To quantify how Ca²⁺ cluster size was altered in *Cxcr4*-cKO mice at each developmental stage, we followed a bootstrapping and regression-based approach:
5. From each experimental group (*Cxcr4*-cKO and age-matched controls), we randomly selected 900 Ca²⁺ clusters and sorted them by size.
6. This procedure was repeated 100 times, and the mean cluster size within each rank bin was calculated to generate smooth, sorted size distributions for both *Cxcr4*-cKO and control.
7. When plotting the *Cxcr4*-cKO versus control sorted cluster sizes, the data initially deviated from the identity line but then converged after a “kink.” We identified this kink as the point that maximized the perpendicular distance to the identity line.

Linear Regression Below Kink: For all clusters below this kink we computed the Pearson correlation coefficient between *Cxcr4*-cKO and control sizes and performed a linear regression constrained through the origin. The resulting slope (β) was taken as the empirical scaling factor by which cluster size was increased in *Cxcr4*-cKO relative to control.

#### Statistical analysis of Ca^2+^ imaging data

Statistical analyses were performed using MATLAB (2023a), OriginPro (2025) and IBM SPSS Statistics (version 28). Population data are reported as mean ± standard error of the mean (SEM), unless stated otherwise. Multi-group data were analyzed using generalized linear models, with either a normal distribution (identity link) or gamma distribution (log link), based on a priori considerations and model fit, as assessed using the small-sample corrected Akaike Information Criterion. The fixed effects included ‘genotype’, ‘age group’, and their interaction. Overall model significance was evaluated using the likelihood ratio chi-square test. Fixed effects were examined only if the overall model was statistically significant. When a significant interaction was found, simple contrasts were used to assess the effect of one factor at each level of the other. To prevent overfitting, non-significant interaction terms were removed through hierarchical model selection, provided that Akaike Information Criterion indicated improved fit relative to the saturated model.

For unpaired group comparison the following statistical strategy was applied. The Shapiro–Wilk test was used to test for normality of the data. Parametric testing procedures were applied for normally distributed data; otherwise non-parametric tests were used. In the case of two-sample t-tests and unequal group variances, Welch’s correction was applied. p values (two-tailed tests) <0.05 were considered statistically significant.

## Acknowledgments

This study was supported by the Deutsche Forschungsgemeinschaft (STU 295/10-1; HO 2156/6-1; KI 1816/7-1 #448069679) to R.S, K.H, and K.K, IZKF Jena research grant (#J53) to P.A., and Germany’s Excellence Strategy – EXC2151 – 390873048 (to N.B-S.). We thank Christine Anders, Ina Ingrisch, Heike Stadler, and the ZFW of UKJ for technical assistance and Elvira Mass (LIMES Institute Bonn) for assistance with bulk-RNA-Seq. Drawings in Figures 2F, 3A, and S4B were derived from scidraw.io (CC-BY license) and were modified – we thank the creators for sharing.

## CRediT authorship contribution statement in alphabetical order

Conceptualization, P.A., K.H., K.K., and R.S.;

Data curation, P.A., J.G., R.S., and S.V.;

Formal analysis, P.A., J.G., H.H., K.K., N.B-S., R.S., and S.V.;

Funding acquisition, P.A., K.H., K.K., and R.S.;

Investigation, P.A., P.A-K., J.G., H.H., C.M., D.R., O.S., R.S., and S.V.;

Methodology, P.A., P.A-K., J.G., H.H., K.K., and R.S.;

Project administration, P.A., and R.S.;

Resources, P.A., K.H., K.K., C.M., D.S., and R.S.;

Software, J.G., K.K.;

Supervision, P.A., K.H., K.K., C.M., and R.S.;

Visualization, P.A., J.G., K.K, and R.S.;

Writing – original draft, P.A., J.G., K.K., and R.S.;

Writing – review and editing, P.A., J.G., K.H., K.K., and R.S.

---

**Figure S1 (relates to Figure 1).**
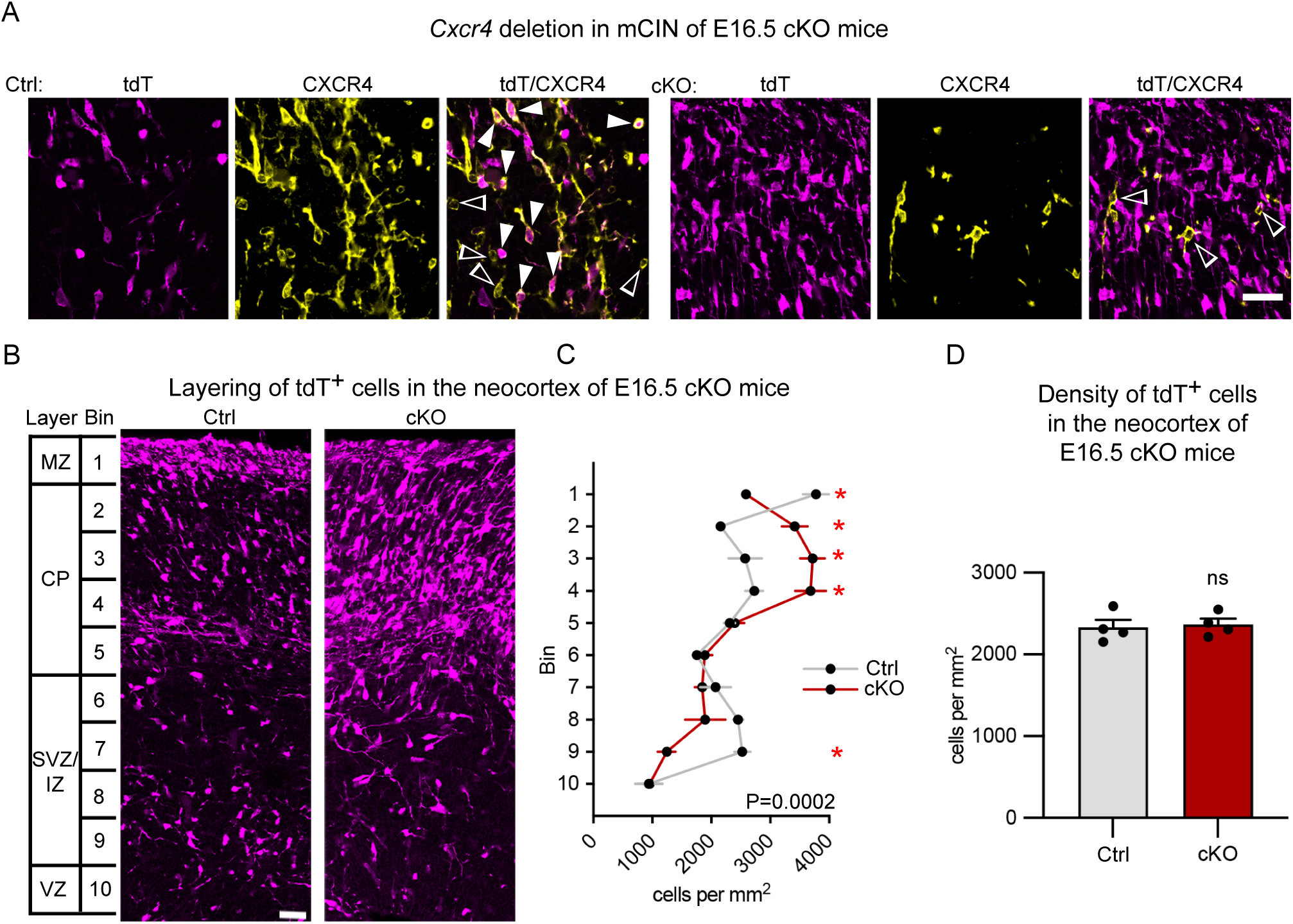
Efficient *Cxcr4* knockout in *Cxcr4*-cKOs. **A,** Confocal images demonstrate immunofluorescence for tdT and CXCR4 in the neocortex of E16.5 control and *Cxcr4*-cKO mice (*Nkx2.1*-Cre; *R26*^LSL-tdT^). Numerous tdT^+^ CXCR4^+^ cells (filled arrowheads) are present in control but not in *Cxcr4*-cKO mice whereas CXCR4^+^ tdT^-^ cells (empty arrowheads) exist in both conditions. **B**, Confocal images demonstrate abnormal layering of tdT^+^ cells in the neocortex of *Cxcr4*-cKO mice. **C,** Line graph showing the distribution of tdT^+^ cells in the neocortex as mean density per bin +/- SEM. Cells were assigned to one of ten bins according to their relative position between pia mater and ventricle (see ***B*** for bins and cortical layers). p = 0.0002 for bin-genotype-interaction as by two-way ANOVA with repeated measures; *: false discovery rate (q) <0.1 as by two-stage linear step-up procedure. **D**, Bar graph shows the density of tdT^+^ cells in the neocortex (averaged across all layers). Circles represent individual mice. **Abbreviations:** cKO, *Cxcr4*-cKO; CP, cortical plate; Ctrl, control; IZ, intermediate zone; MZ, marginal zone; ns; not significant; SVZ; subventricular zone. **Scale bars:** 30 µm in ***A*** and ***B***.

**Figure S2 (relates to Figure 1).**
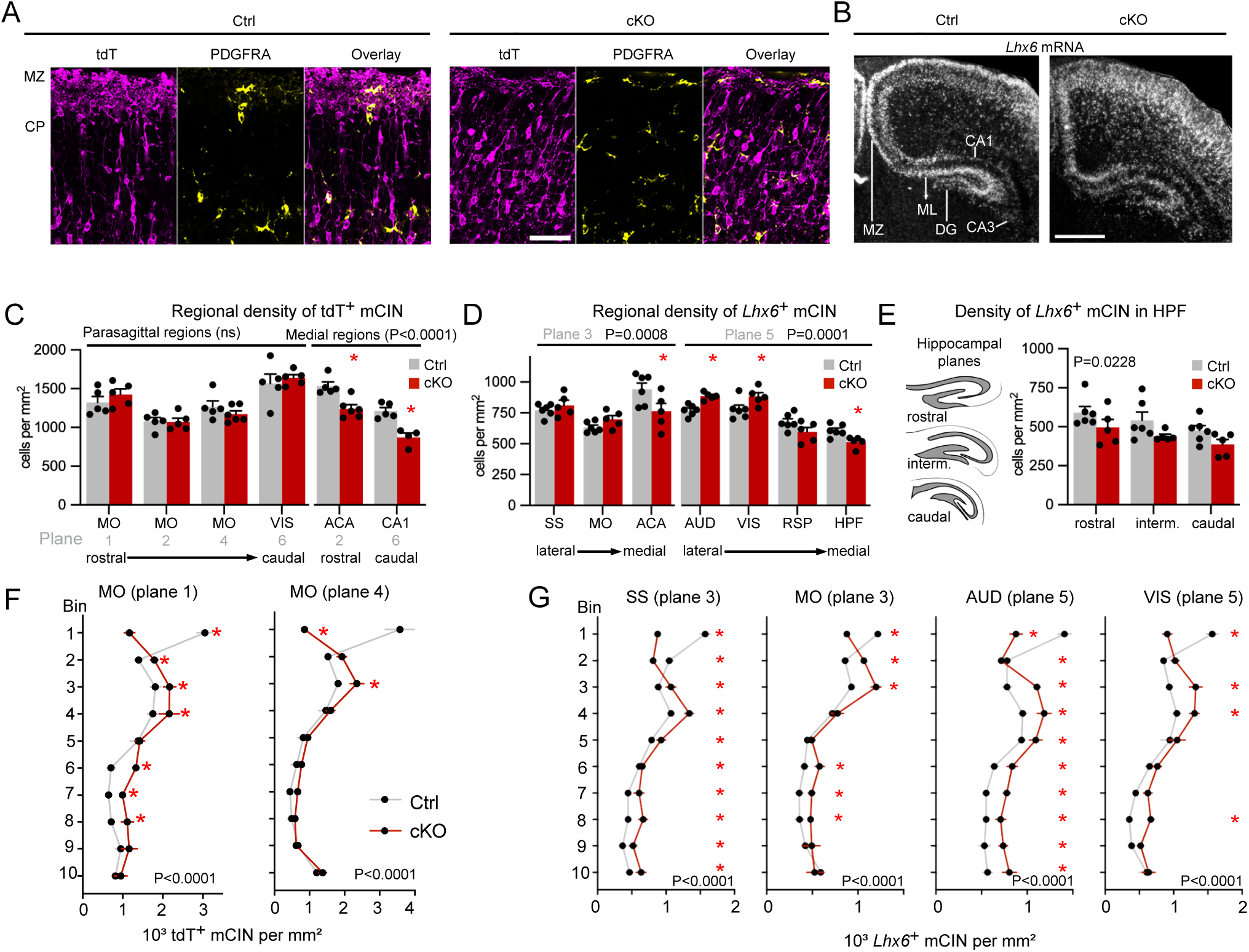
Regional distribution and layering of mCIN in P0 *Cxcr4*-cKO mice. **A**, Confocal images demonstrate immunofluorescence for tdT and PDGFRA in the neocortical MZ and subjacent cortical plate of P0 control and *Cxcr4*-cKO mice. **B,** Darkfield micrographs of hybridized frontal sections (plane 5) demonstrate *Lhx6* in the neocortex and hippocampus. **C-E**, Bar graphs show the density of PDGFRA^-^ tdT^+^ (***C***) and *Lhx6*^+^ (***D***, ***E***) mCIN for the indicated cortical regions and sectional planes in cells per mm^2^ (sectional planes and regions are illustrated in Figures 1A and **1B**). Data shown in **C** and **D** were used to generate graphs shown in Figures 1A and **1B**. **F,G,** Line graphs show the distribution of PDGFRA^-^ tdT^+^ mCIN (***F***) and *Lhx6*^+^ mCIN (***G***) between MZ and VZ in the indicated cortical areas. See Figure 1C for layers and bins. **Statistics:** p-values for genotype-bin interaction as by two-way ANOVA with repeated measures; *: false discovery rate (q) <0.1 as by two-stage linear step-up procedure (***C–G***). **Cortical regions:** ACA, anterior cingulate; AUD, auditory; CA, *Cornu ammonis*, HPF, hippocampal formation; MO, motor; RSP, retrosplenial; SS, somatosensory; VIS, visual; DG, dentate gyrus. **Abbreviations:** cKO, *Cxcr4*-cKO; CP, cortical plate; Ctrl, control; interm., intermediate; ML, molecular layer; MZ, marginal zone; ns not significant. All graphs show mean values + SEM. Circles in ***C* - *E*** represent individual mice. **Scale bars:** 50 µm in ***A***, 500 µm in ***B***.

**Figure S3.**
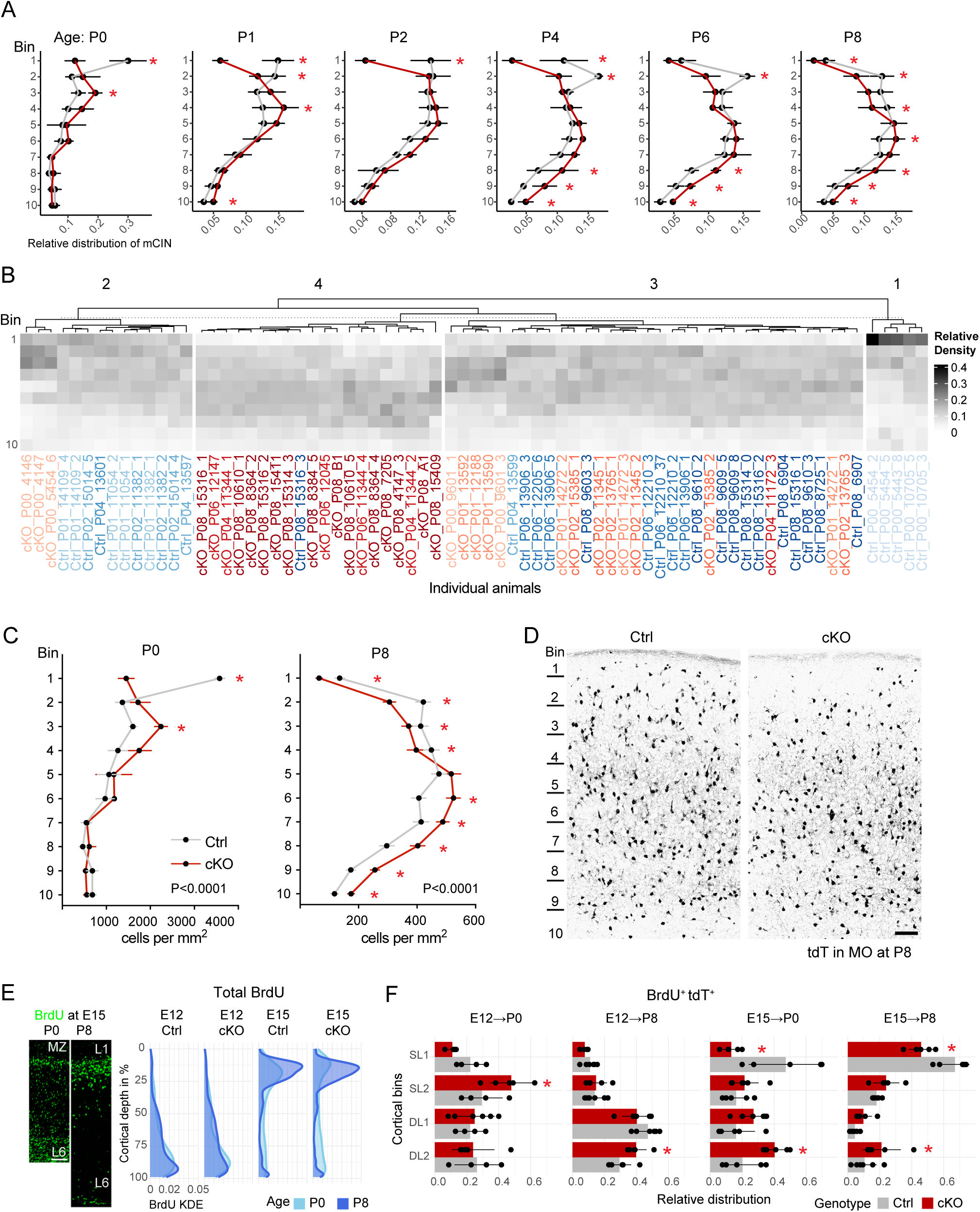
(relates to Figure 2). Postnatal layer allocation of mCIN in *Cxcr4*-cKO mice. **A,** Line graphs show the distribution of mCIN across layers 1 to 6 of the motor cortex at the indicated postnatal stages as relative density. Data in ***A*** refer to Figure 2C. **B,** Heatmaps show the distribution of mCIN across ten cortical bins as relative density. Each analyzed animal is depicted and identified by label. The Clustering of the distribution highlights the differences between ages and genotypes. Data in ***B*** refer to Figure 2E. **C,** Line graphs show the distribution of mCIN across layers 1 to 6 of the motor cortex on P0 and P8. **D,** Representative images demonstrate tdT in the motor cortex on P8. **A - C**, Data represent PDGFRA tdT^+^ cells (i.e. mCIN). **E, Left,** Confocal images demonstrate the distribution of E15-pulsed BrdU^+^ cells in the motor cortex at P0 and P8. Note that E15-pulsed cells accumulate beneath the MZ between P0 and P8. **Right,** Density plots show the distribution of BrdU^+^ cells (i.e. non-mCIN and mCIN) across layers 1 to 6 of the neocortex. BrdU was applied on E12 or E15 and animals assessed on P0 or P8 as indicated. The position within the cortex is represented as relative cortical depth on the y-axis. Density is shown as kernel density estimates (KDE) on the x-axis. **F,** Bar graphs show the distribution of E12-pulsed and E15-pulsed BrdU^+^ tdT^+^ cells (i.e. mCIN and some first-wave oligodendrocytes) in the neocortex on P0 and P8 as relative density. The cortex was divided into four bins for superficial and deep layers (SL1, SL2, DL1, DL2). **Statistics A,F,** *: p <0.1 as by beta regression model followed by simultaneous tests for general linear hypotheses for repeated measurements correction. **C,** p-values for genotype-bin interaction as by two-way ANOVA with repeated measures; *: false discovery rate (q) <0.1 as by two-stage linear step-up procedure. All graphs show mean + SEM. Source data are provided in **Table S5**. **Scale bar:** 100 µm in ***D***. **Abbreviations:** cKO, *Cxcr4*-cKO; Ctrl, control; DL and SL, deep and superficial cortical layers; L1 and L6, cortical layers 1 and 6; MZ, marginal zone.

**Figure S4 (relates to Figure 3).**
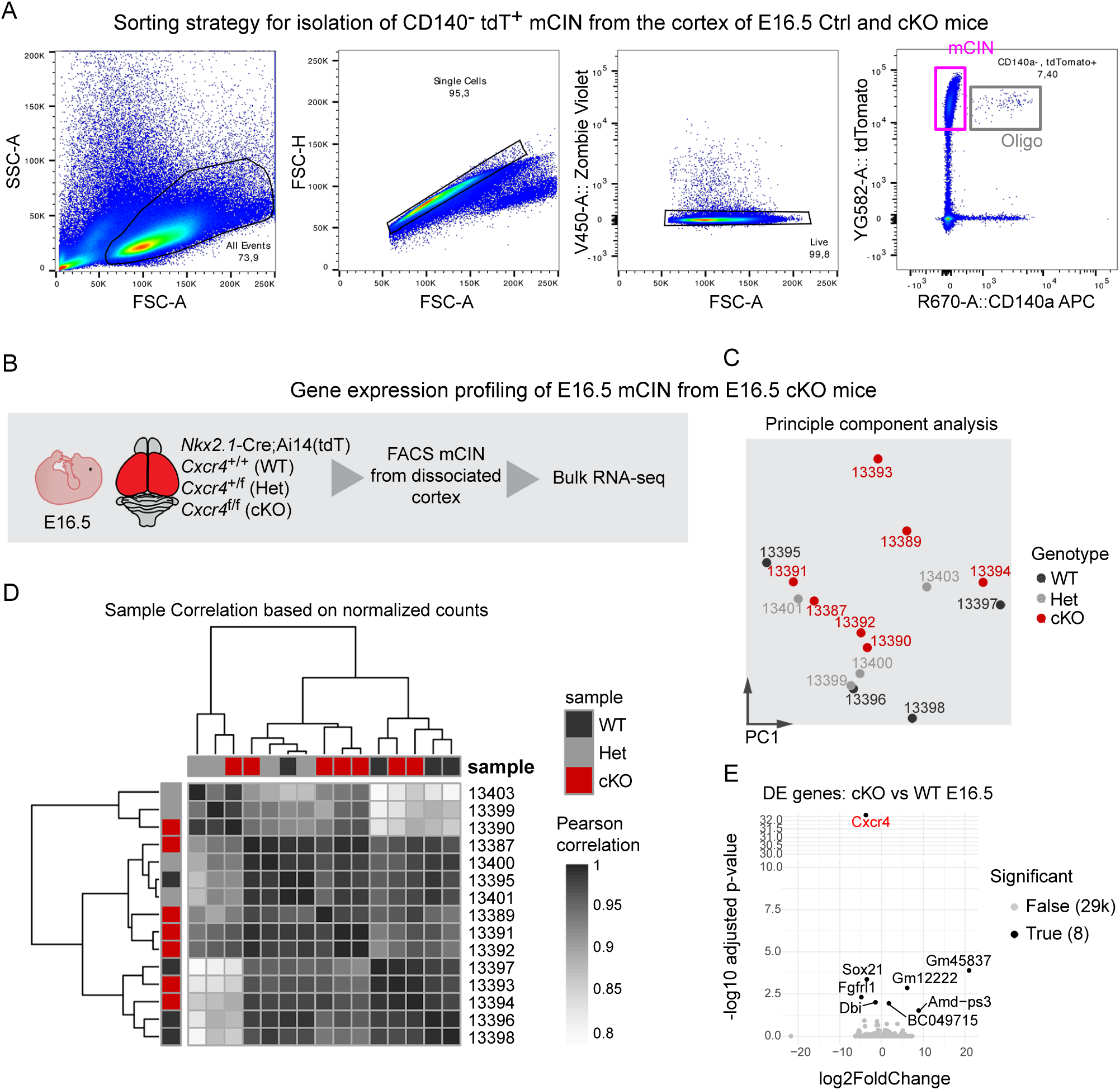
Unchanged gene expression profiles of ectopically migrating E16.5 mCIN. **A**, Scatter plots demonstrate the sorting strategy used to isolate mCIN from the cortex of E16.5 control and *Cxcr4*-cKO mice. The sorting gate was placed on single live CD140^-^ tdT^+^ cells. First wave oligodendrocytes (Oligo) are CD140^+^ tdT^+^ and contribute < than 1% to the tdT^+^ population **B,** Experimental procedure used for gene expression profiling of E16.5 *Cxcr4*-deficient mCIN. **C,D,** Principle component analysis with all present genes (***C***) and pearson correlation followed by hierarchical clustering based on normalized counts (***D***) show no clear segregation of samples from *Cxcr4*-wildtype (WT), *Cxcr4*-heterozygous (Het), and *Cxcr4*-cKO (cKO) mice. ***E,*** The Volcano plot shows 8 differentially expressed (DE) genes out of 29.000 genes in *Cxcr4*-deficient mCIN (*Cxcr4*-cKO versus wildtype).

**Figure S5 (relates to Figure 3).**
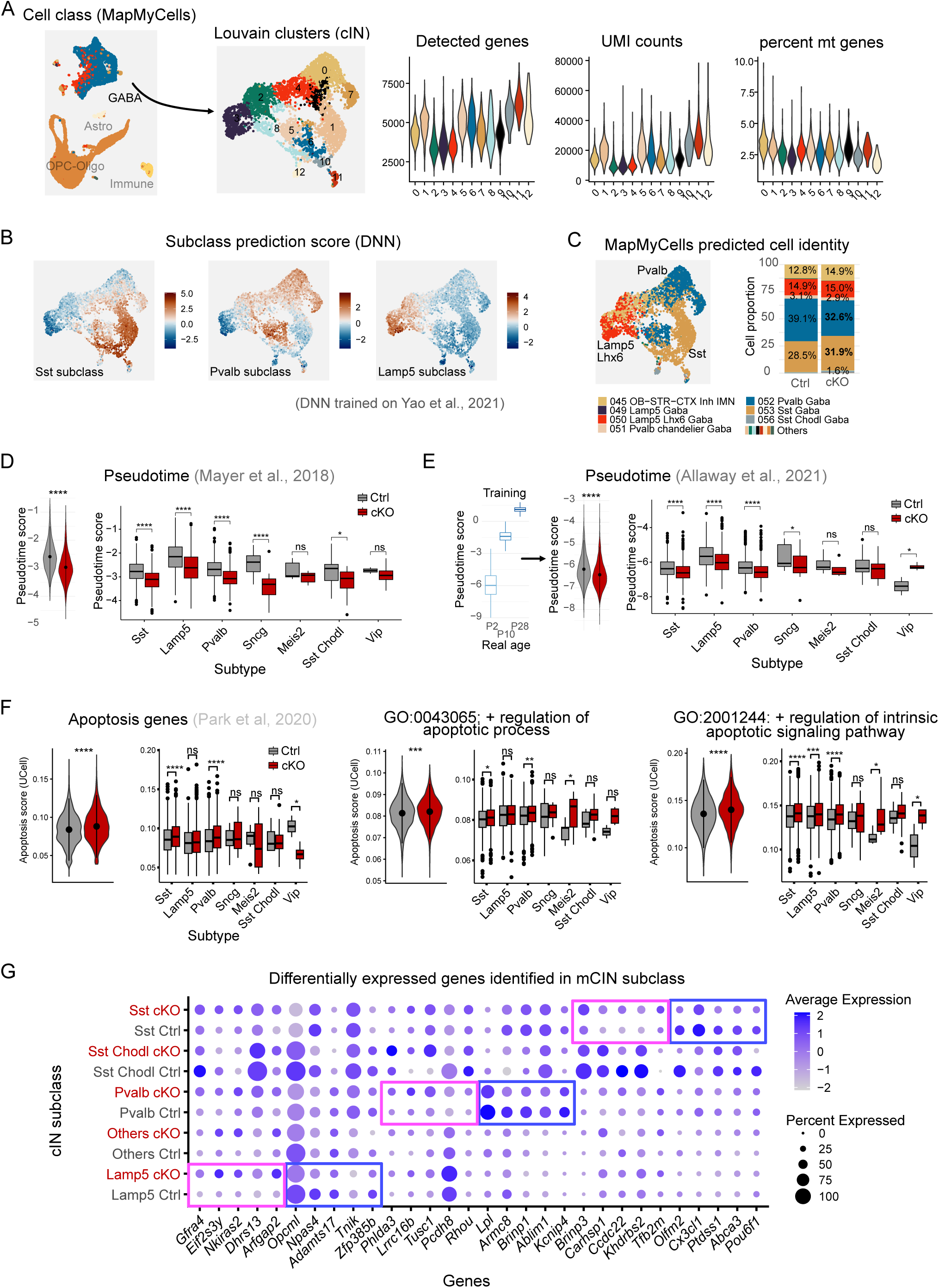
Single-cell RNA sequencing quality control, prediction, and pseudotime analysis of mCIN. **A,** Pipeline for selecting quality-controlled mCIN. Violin plots display the distribution of key quality control parameters (number of genes per cell, UMI counts, and percentage of mitochondrial [mt] genes) across each Louvain cluster. **B,** The heatmap shows the Deep Neural Network (DNN)-derived prediction scores for *Sst*, *Pvalb*, and *Lamp5* subclasses used for the assignment in Figure 3C (red indicates high prediction score, blue indicates low). **C,** Cell-type prediction performed using the online tool MapMyCells. Consistent with the findings in Figure 3, the proportion of the *Lamp5 Lhx6* subclass was similar in both genotypes, but the ratio of the *Sst* and *Pvalb* subclasses was altered in *Cxcr4*-cKOs. **D,** Pseudotime scores derived from applying the ordinal regression model trained on dataset from ^37^ for all mCIN (left) and per subclass (right) in control and *Cxcr4*-cKOs. **E, Left,** Training of a pseudotime ordinal regression model on cells from P2, P10, and P28 animals ^49^. **Center,** The resulting pseudotime model was then used to predict the age of all mCIN in control and *Cxcr4*-cKO. **Right,** Predicted age distribution per subclass in control and *Cxcr4*-cKO. **F,** Apoptosis scoring based on UCell metagene analysis. Scores were calculated using gene sets derived from ^81^ (n=30 genes), GO:0043065 (n=625 genes), and GO:2001244 (n=73 genes) for all mCIN and subclasses. **G,** Dot plot highlighting top five genes within major *Sst*, *Pvalb*, and *Lamp5* subclasses for control and *Cxcr4*-cKO. Refers also to **Tables S2** and **S5** and Figure 3G. Genes were selected on adjusted p value <0.05 and percentage expression >0.3. **Statistics**: Two-group comparisons were analyzed using the Kolmogorov-Smirnov test. For subclass comparisons, a multiple comparison of means was performed using the *emmeans* function with Bonferroni p-value adjustment (***D*** - ***F***).

**Figure S6 (relates to Figure 4).**
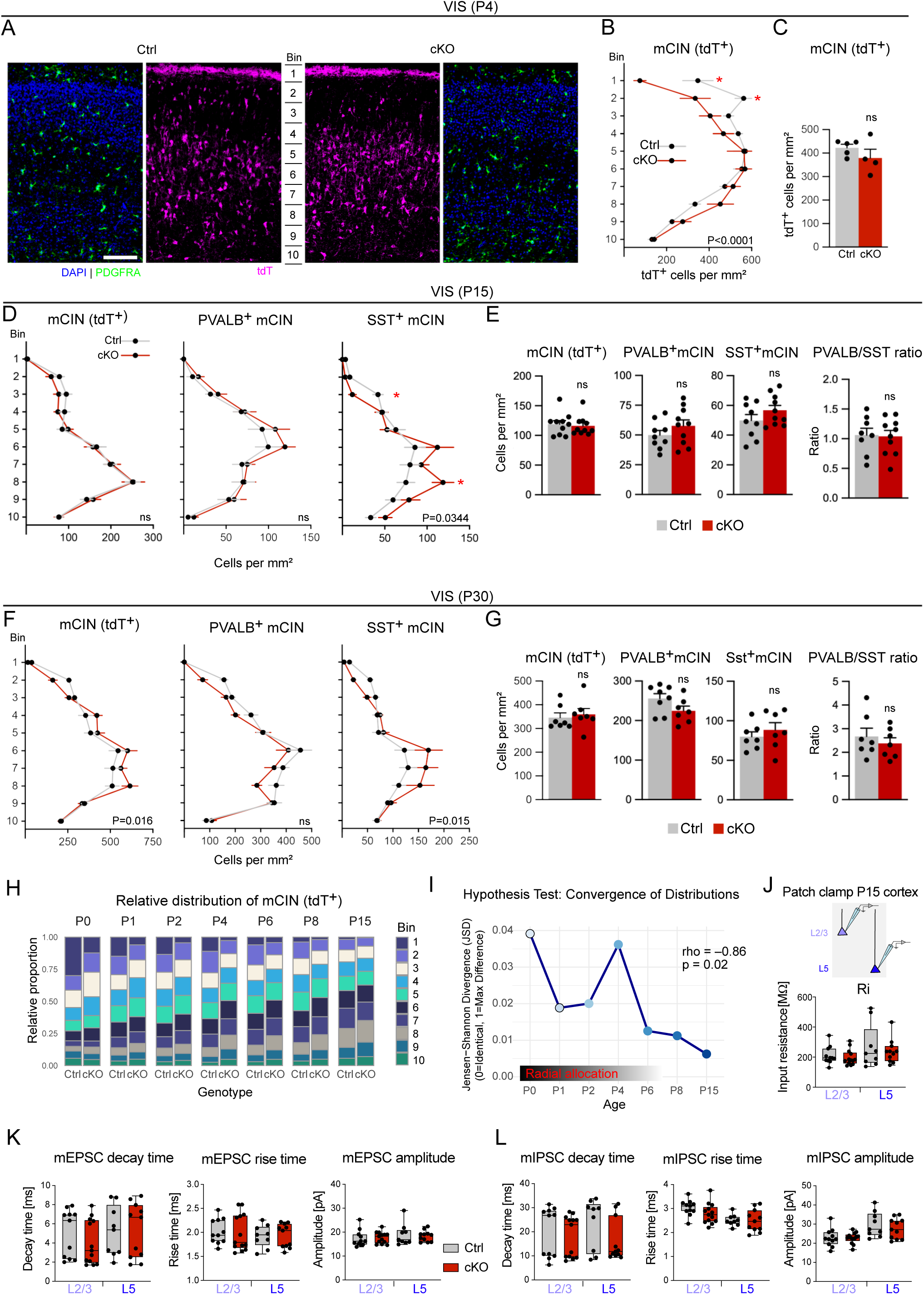
Downward shift of SST^+^ mCIN at the mature stage. **A,** Confocal images demonstrate tdT and PDGFRA immunofluorescences in the visual cortex (VIS) of P4 control and *Cxcr4*-cKO mice (*Nkx2.1*-Cre; *Rosa26*^LSL-tdT^). **B-G**, Line and bar graphs show the distribution and total number of tdT^+^, PVALB^+^, and SST^+^ mCIN across layers 1 to 6 of the VIS at P4 ***(B,C)***, P15 ***(D,E),*** and P30 ***(F,G)***. PVALB/SST ratios are shown in ***E*** and ***G*** to the right. Graphs show mean + SEM. Circles represent individual mice (***C,E,G***). **H**, Proportion of each bin in the total mCIN count shown for the indicated timepoints (same data as in Figures 2C**, 4A,** and **S3A**). **I,** Jensen-Shannon divergence calculated on the distribution of control and *Cxcr4*-cKO mCIN (tdT^+^) across time (0∼ identical, 1∼max difference). The decreasing trend (negative correlation: rho= 0.86) suggests a convergence of genotype distributions with maturation. **J-L,** Miniature excitatory and miniature inhibitory postsynaptic potentials (mEPSC, mIPSC) recorded in pyramidal cells in layer 2/ 3 (L2/3) and layer 5 (L5) of the VIS at P15 – P18. Box and whisker plots (minimum to maximum with median represented as horizontal line and individual cells as circles) show the input resistande (R_i_), decay and rise time, and amplitude. **Statistics: B,D,F,** p-values for genotype-bin-interaction as by two-way ANOVA with repeated measures; *false discovery rate (q) <0.1 as by two-stage linear step-up procedure. **C,E,G,J-L,** No significant differences as by Mann-Whitney test. **Abbreviations:** cKO, *Cxcr4*-cKO; Ctrl, control; ns, not significant. **Scale bar:** 100 µm in ***A***.

**Figure S7 (relates to Figure 4).**
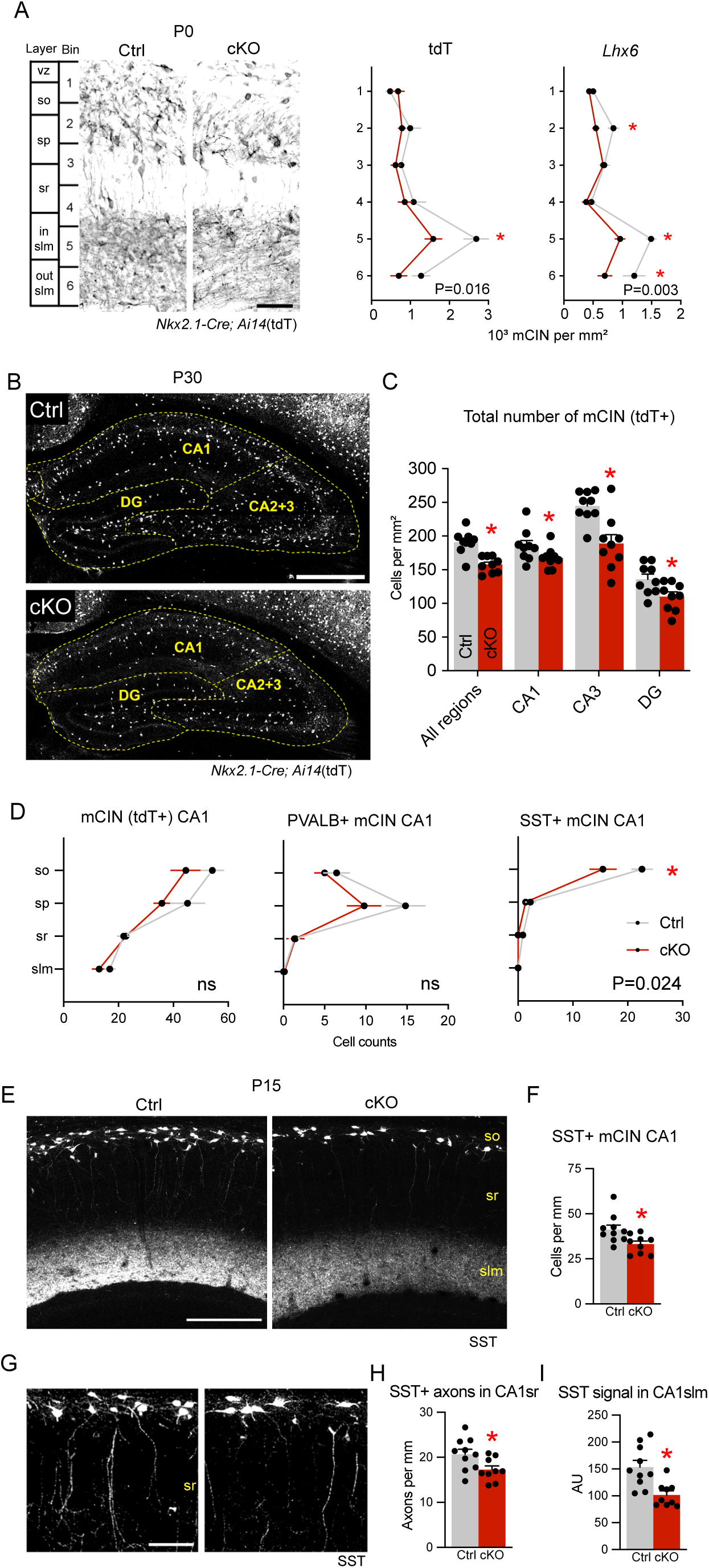
Perturbed TM reduces the overall mCIN density in the hippocampus without affecting laminar distribution. **A,** Distribution of mCIN (PDGFR^-^ tdT^+^) in the hippocampal CA1 region at P0. mCIN were detected and quantified across six bins by using *Nkx2.1-Cre* lineage marker tdT (micrograph and left line graph) and *Lhx6* mRNA (right line graph). **B**, tdT immunofluorescence in the hippocampus at P30 in control and *Cxcr4*-cKO mice. **C**, Bar graphs show the density of mCIN for the hippocampus and hippocampal subregions CA1, CA3, and DG. **D**, Line graphs show the distribution of tdT^+^, PVALB ^+^, and SST^+^ mCIN across the hippocampal layers in the CA1 at P30 as cell count per field of view. **E,G**, Micrographs demonstrate SST immunofluorescence in the CA1 at P15 at low (***E***) and high magnification (***G***). **F,H,I,** Quantification of SST^+^ mCIN in *stratum oriens* (***F***), SST^+^ axons in *stratum radiatum* (***H***), and SST-signal in *stratum lacunosum moleculare* in CA1 at P15. **Statistics**: **A,C,D,** p-values for genotype (***A***, tdT; ***C***) and layer-genotype interaction (***A***, *Lhx6*; ***D***) as by two-way ANOVA with repeated measures; *: false discovery rate (q) <0.1 as by two-stage linear step-up procedure. ***F,H,I,*** p-values as by Mann-Whitney test. ***A-I,*** All data are mean +/- SEM. Source data are provided in **Table S5**. **Abbreviations**: CA, *cornu ammonis*; cc, *corpus callosum*; cKO, *Cxcr4*-cKO; Ctrl, control; DG, dentate gyrus; slm, *stratum lacunosum moleculare*;in slm and out slm, inner and outer *slm*; sp, pyramidal cell layer; so, *stratum oriens*; sr, *stratum radiatum*; VZ ventricular zone. **Scale bars:** 45 µm in ***A***, 500 µm in ***B***, 630 µm in ***E,*** 100 µm in ***G***.

**Figure S8 (relates to Figure 6).**
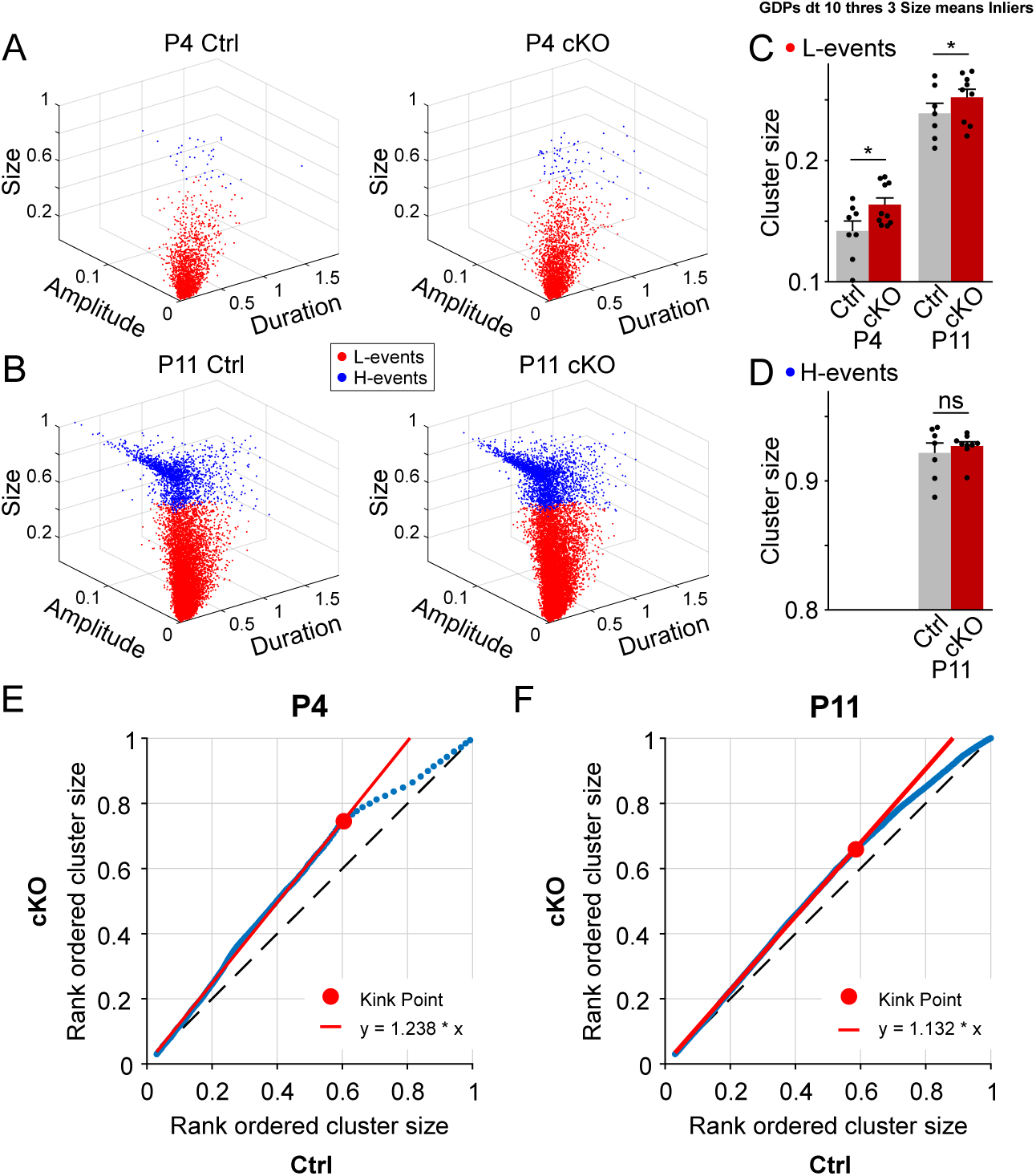
Increased spatial extent of L-events in P4 and P11 *Cxcr4*-cKO mice. **A,B,** 3D plots of all detected Ca^2+^ clusters in control and *Cxcr4*-cKO mice, illustrating their size, amplitude, and duration. L-events were identified in P4 control mice based on the 98th percentile to define margins within the 3D scatter plot. Data points falling outside these margins were classified as H-events. The same thresholds were then applied to the corresponding scatter plots in P4 *Cxcr4*-cKO and P11 datasets. **C,** L-event size was significantly increased in *Cxcr4*-cKO mice at both P4 and P11. **D,** No significant differences were found for H-events at P11. Due to the low number of H-events at P4, no statistical testing was performed for this group. **E,** Rank-ordered plot of Ca²⁺ cluster size in *Cxcr4*-cKO versus control mice at P4. The red line represents the linear regression of all data points to the left of the kink point (i.e., largely corresponding to L-events). **F,** Same as in ***E***, but for the P11 dataset. **A-B, E-F,** Each data point represents a single Ca^2+^ cluster. ***C,D,*** Each circle represents an individual animal. Data are presented as mean ± SEM. *p <0.05.

## References

1. Luhmann HJ. Neurophysiology of the Developing Cerebral Cortex: What We Have Learned and What We Need to Know. Front Cell Neurosci. 2021;15:814012.

2. Le Magueresse C, Monyer H. GABAergic interneurons shape the functional maturation of the cortex. Neuron. 2013;77(3):388–405.

3. Anderson SA, Eisenstat DD, Shi L, Rubenstein JL. Interneuron migration from basal forebrain to neocortex: dependence on Dlx genes. Science. 1997;278(5337):474–6.

4. Marin O. Development of GABAergic Interneurons in the Human Cerebral Cortex. Eur J Neurosci. 2025;61(9):e70136.

5. Contractor A, Ethell IM, Portera-Cailliau C. Cortical interneurons in autism. Nat Neurosci. 2021;24(12):1648–59.

6. Marin O. Parvalbumin interneuron deficits in schizophrenia. Eur Neuropsychopharmacol. 2024;82:44–52.

7. Parnell AA, McCarthy DM, Trupiano MX, Schatschneider C, Bhide PG. Nicotine in e-cigarette aerosol reduces GABA neuron migration via the alpha7 nicotinic acetylcholine receptor. Sci Rep. 2025;15(1):35635.

8. Schmidt MJ, Mirnics K. Neurodevelopment, GABA system dysfunction, and schizophrenia. Neuropsychopharmacology. 2015;40(1):190–206.

9. Xu Q, Tam M, Anderson SA. Fate mapping Nkx2.1-lineage cells in the mouse telencephalon. J Comp Neurol. 2008;506(1):16–29.

10. Huang ZJ, Paul A. The diversity of GABAergic neurons and neural communication elements. Nat Rev Neurosci. 2019;20(9):563–72.

11. Lim L, Mi D, Llorca A, Marin O. Development and Functional Diversification of Cortical Interneurons. Neuron. 2018;100(2):294–313.

12. Lavdas AA, Grigoriou M, Pachnis V, Parnavelas JG. The medial ganglionic eminence gives rise to a population of early neurons in the developing cerebral cortex. J Neurosci. 1999;19(18):7881–8.

13. Lopez-Bendito G, Sanchez-Alcaniz JA, Pla R, Borrell V, Pico E, Valdeolmillos M, et al. Chemokine signaling controls intracortical migration and final distribution of GABAergic interneurons. J Neurosci. 2008;28(7):1613–24.

14. Tanaka DH, Yanagida M, Zhu Y, Mikami S, Nagasawa T, Miyazaki J, et al. Random walk behavior of migrating cortical interneurons in the marginal zone: time-lapse analysis in flat-mount cortex. J Neurosci. 2009;29(5):1300–11.

15. Tanaka DH, Maekawa K, Yanagawa Y, Obata K, Murakami F. Multidirectional and multizonal tangential migration of GABAergic interneurons in the developing cerebral cortex. Development. 2006;133(11):2167–76.

16. Danglot L, Triller A, Marty S. The development of hippocampal interneurons in rodents. Hippocampus. 2006;16(12):1032–60.

17. Toudji I, Toumi A, Chamberland E, Rossignol E. Interneuron odyssey: molecular mechanisms of tangential migration. Front Neural Circuits. 2023;17:1256455.

18. Stumm RK, Zhou C, Ara T, Lazarini F, Dubois-Dalcq M, Nagasawa T, et al. CXCR4 regulates interneuron migration in the developing neocortex. J Neurosci. 2003;23(12):5123–30.

19. Bartolini G, Sanchez-Alcaniz JA, Osorio C, Valiente M, Garcia-Frigola C, Marin O. Neuregulin 3 Mediates Cortical Plate Invasion and Laminar Allocation of GABAergic Interneurons. Cell Rep. 2017;18(5):1157–70.

20. Wang Y, Li G, Stanco A, Long JE, Crawford D, Potter GB, et al. CXCR4 and CXCR7 have distinct functions in regulating interneuron migration. Neuron. 2011;69(1):61–76.

21. Sanchez-Alcaniz JA, Haege S, Mueller W, Pla R, Mackay F, Schulz S, et al. Cxcr7 controls neuronal migration by regulating chemokine responsiveness. Neuron. 2011;69(1):77–90.

22. Abe P, Molnar Z, Tzeng YS, Lai DM, Arnold SJ, Stumm R. Intermediate Progenitors Facilitate Intracortical Progression of Thalamocortical Axons and Interneurons through CXCL12 Chemokine Signaling. J Neurosci. 2015;35(38):13053–63.

23. Abe P, Mueller W, Schutz D, MacKay F, Thelen M, Zhang P, et al. CXCR7 prevents excessive CXCL12-mediated downregulation of CXCR4 in migrating cortical interneurons. Development. 2014;141(9):1857–63.

24. Li G, Adesnik H, Li J, Long J, Nicoll RA, Rubenstein JL, et al. Regional distribution of cortical interneurons and development of inhibitory tone are regulated by Cxcl12/Cxcr4 signaling. J Neurosci. 2008;28(5):1085–98.

25. Saaber F, Schutz D, Miess E, Abe P, Desikan S, Ashok Kumar P, et al. ACKR3 Regulation of Neuronal Migration Requires ACKR3 Phosphorylation, but Not beta-Arrestin. Cell Rep. 2019;26(6):1473–88 e9.

26. Volk DW, Chitrapu A, Edelson JR, Lewis DA. Chemokine receptors and cortical interneuron dysfunction in schizophrenia. Schizophr Res. 2015;167(1-3):12–7.

27. Eid L, Lokmane L, Raju PK, Tene Tadoum SB, Jiang X, Toulouse K, et al. Both GEF domains of the autism and developmental epileptic encephalopathy-associated Trio protein are required for proper tangential migration of GABAergic interneurons. Mol Psychiatry. 2025;30(4):1338–58.

28. Sun X, Wang L, Wei C, Sun M, Li Q, Meng H, et al. Dysfunction of Trio GEF1 involves in excitatory/inhibitory imbalance and autism-like behaviors through regulation of interneuron migration. Mol Psychiatry. 2021;26(12):7621–40.

29. Toritsuka M, Kimoto S, Muraki K, Landek-Salgado MA, Yoshida A, Yamamoto N, et al. Deficits in microRNA-mediated Cxcr4/Cxcl12 signaling in neurodevelopmental deficits in a 22q11 deletion syndrome mouse model. Proc Natl Acad Sci U S A. 2013;110(43):17552–7.

30. Meechan DW, Tucker ES, Maynard TM, LaMantia AS. Cxcr4 regulation of interneuron migration is disrupted in 22q11.2 deletion syndrome. Proc Natl Acad Sci U S A. 2012;109(45):18601–6.

31. Cash-Padgett T, Sawa A, Jaaro-Peled H. Increased stereotypy in conditional Cxcr4 knockout mice. Neurosci Res. 2016;105:75–9.

32. Mayer C, Hafemeister C, Bandler RC, Machold R, Batista Brito R, Jaglin X, et al. Developmental diversification of cortical inhibitory interneurons. Nature. 2018;555(7697):457–62.

33. Vogt D, Hunt RF, Mandal S, Sandberg M, Silberberg SN, Nagasawa T, et al. Lhx6 directly regulates Arx and CXCR7 to determine cortical interneuron fate and laminar position. Neuron. 2014;82(2):350–64.

34. Tanaka DH, Mikami S, Nagasawa T, Miyazaki J, Nakajima K, Murakami F. CXCR4 is required for proper regional and laminar distribution of cortical somatostatin-, calretinin-, and neuropeptide Y-expressing GABAergic interneurons. Cereb Cortex. 2010;20(12):2810–7.

35. Schonemeier B, Kolodziej A, Schulz S, Jacobs S, Hoellt V, Stumm R. Regional and cellular localization of the CXCl12/SDF-1 chemokine receptor CXCR7 in the developing and adult rat brain. J Comp Neurol. 2008;510(2):207–20.

36. Venkataramanappa S, Saaber F, Abe P, Schutz D, Kumar PA, Stumm R. Cxcr4 and Ackr3 regulate allocation of caudal ganglionic eminence-derived interneurons to superficial cortical layers. Cell Rep. 2022;40(5):111157.

37. Stumm R, Kolodziej A, Schulz S, Kohtz JD, Hollt V. Patterns of SDF-1alpha and SDF-1gamma mRNAs, migration pathways, and phenotypes of CXCR4-expressing neurons in the developing rat telencephalon. J Comp Neurol. 2007;502(3):382–99.

38. Lysko DE, Putt M, Golden JA. SDF1 regulates leading process branching and speed of migrating interneurons. J Neurosci. 2011;31(5):1739–45.

39. Atkins M, Wurmser M, Darmon M, Roche F, Nicol X, Metin C. CXCL12 targets the primary cilium cAMP/cGMP ratio to regulate cell polarity during migration. Nat Commun. 2023;14(1):8003.

40. Lysko DE, Putt M, Golden JA. SDF1 reduces interneuron leading process branching through dual regulation of actin and microtubules. J Neurosci. 2014;34(14):4941–62.

41. Kessaris N, Fogarty M, Iannarelli P, Grist M, Wegner M, Richardson WD. Competing waves of oligodendrocytes in the forebrain and postnatal elimination of an embryonic lineage. Nat Neurosci. 2006;9(2):173–9.

42. Hevner RF, Daza RA, Englund C, Kohtz J, Fink A. Postnatal shifts of interneuron position in the neocortex of normal and reeler mice: evidence for inward radial migration. Neuroscience. 2004;124(3):605–18.

43. Yao Z, van Velthoven CTJ, Nguyen TN, Goldy J, Sedeno-Cortes AE, Baftizadeh F, et al. A taxonomy of transcriptomic cell types across the isocortex and hippocampal formation. Cell. 2021;184(12):3222–41 e26.

44. Abe P, Lavalley A, Morassut I, Santinha AJ, Roig-Puiggros S, Javed A, et al. Molecular programs guiding arealization of descending cortical pathways. Nature. 2024;634(8034):644–51.

45. Allaway KC, Gabitto MI, Wapinski O, Saldi G, Wang CY, Bandler RC, et al. Genetic and epigenetic coordination of cortical interneuron development. Nature. 2021;597(7878):693–7.

46. Southwell DG, Paredes MF, Galvao RP, Jones DL, Froemke RC, Sebe JY, et al. Intrinsically determined cell death of developing cortical interneurons. Nature. 2012;491(7422):109–13.

47. Wong FK, Bercsenyi K, Sreenivasan V, Portales A, Fernandez-Otero M, Marin O. Pyramidal cell regulation of interneuron survival sculpts cortical networks. Nature. 2018;557(7707):668–73.

48. Zechel S, Fernandez-Suarez D, Ibanez CF. Cell-autonomous role of GFRalpha1 in the development of olfactory bulb GABAergic interneurons. Biol Open. 2018;7(5).

49. Yang J, Serrano P, Yin X, Sun X, Lin Y, Chen SX. Functionally distinct NPAS4-expressing somatostatin interneuron ensembles critical for motor skill learning. Neuron. 2022;110(20):3339–55 e8.

50. Shepard R, Heslin K, Hagerdorn P, Coutellier L. Downregulation of Npas4 in parvalbumin interneurons and cognitive deficits after neonatal NMDA receptor blockade: relevance for schizophrenia. Transl Psychiatry. 2019;9(1):99.

51. Voronova A, Yuzwa SA, Wang BS, Zahr S, Syal C, Wang J, et al. Migrating Interneurons Secrete Fractalkine to Promote Oligodendrocyte Formation in the Developing Mammalian Brain. Neuron. 2017;94(3):500–16 e9.

52. Zhang Z, Ye M, Li Q, You Y, Yu H, Ma Y, et al. The Schizophrenia Susceptibility Gene OPCML Regulates Spine Maturation and Cognitive Behaviors through Eph-Cofilin Signaling. Cell Rep. 2019;29(1):49–61 e7.

53. Talishinsky A, Downar J, Vertes PE, Seidlitz J, Dunlop K, Lynch CJ, et al. Regional gene expression signatures are associated with sex-specific functional connectivity changes in depression. Nat Commun. 2022;13(1):5692.

54. Berkowicz SR, Featherby TJ, Whisstock JC, Bird PI. Mice Lacking Brinp2 or Brinp3, or Both, Exhibit Behaviors Consistent with Neurodevelopmental Disorders. Front Behav Neurosci. 2016;10:196.

55. Kawano H, Nakatani T, Mori T, Ueno S, Fukaya M, Abe A, et al. Identification and characterization of novel developmentally regulated neural-specific proteins, BRINP family. Brain Res Mol Brain Res. 2004;125(1-2):60–75.

56. Ben-Ari Y, Gaiarsa JL, Tyzio R, Khazipov R. GABA: a pioneer transmitter that excites immature neurons and generates primitive oscillations. Physiol Rev. 2007;87(4):1215–84.

57. Kirmse K, Kummer M, Kovalchuk Y, Witte OW, Garaschuk O, Holthoff K. GABA depolarizes immature neurons and inhibits network activity in the neonatal neocortex in vivo. Nat Commun. 2015;6:7750.

58. Kirmse K, Zhang C. Principles of GABAergic signaling in developing cortical network dynamics. Cell Rep. 2022;38(13):110568.

59. Leighton AH, Cheyne JE, Houwen GJ, Maldonado PP, De Winter F, Levelt CN, et al. Somatostatin interneurons restrict cell recruitment to retinally driven spontaneous activity in the developing cortex. Cell Rep. 2021;36(1):109316.

60. Guenthner CJ, Miyamichi K, Yang HH, Heller HC, Luo L. Permanent genetic access to transiently active neurons via TRAP: targeted recombination in active populations. Neuron. 2013;78(5):773–84.

61. Wang Q, Sporns O, Burkhalter A. Network analysis of corticocortical connections reveals ventral and dorsal processing streams in mouse visual cortex. J Neurosci. 2012;32(13):4386–99.

62. Gribizis A, Ge X, Daigle TL, Ackman JB, Zeng H, Lee D, et al. Visual Cortex Gains Independence from Peripheral Drive before Eye Opening. Neuron. 2019;104(4):711–23 e3.

63. Colonnese MT, Kaminska A, Minlebaev M, Milh M, Bloem B, Lescure S, et al. A conserved switch in sensory processing prepares developing neocortex for vision. Neuron. 2010;67(3):480–98.

64. Siegel F, Heimel JA, Peters J, Lohmann C. Peripheral and central inputs shape network dynamics in the developing visual cortex in vivo. Curr Biol. 2012;22(3):253–8.

65. Hinckelmann MV, Dubos A, Artot V, Rudolf G, Nguyen TL, Tilly P, et al. Interneuron migration defects during corticogenesis contribute to Dyrk1a haploinsufficiency syndrome pathogenesis. Mol Psychiatry. 2025;30(11):5227–44.

66. Reichard J, Wolff P, Xie S, Zuo K, Fullio CL, Du J, et al. DNMT1-mediated regulation of somatostatin-positive interneuron migration impacts cortical architecture and function. Nat Commun. 2025;16(1):6834.

67. Wu SJ, Dai M, Yang SP, McCann C, Qiu Y, Kumar V, et al. Pyramidal neurons proportionately alter cortical interneuron subtypes. Nature. 2026;651(8105):421–8.

68. Modol L, Moissidis M, Selten M, Oozeer F, Marin O. Somatostatin interneurons control the timing of developmental desynchronization in cortical networks. Neuron. 2024;112(12):2015–30 e5.

69. Lim L, Pakan JMP, Selten MM, Marques-Smith A, Llorca A, Bae SE, et al. Optimization of interneuron function by direct coupling of cell migration and axonal targeting. Nat Neurosci. 2018;21(7):920–31.

70. Wosniack ME, Kirchner JH, Chao LY, Zabouri N, Lohmann C, Gjorgjieva J. Adaptation of spontaneous activity in the developing visual cortex. Elife. 2021;10.

71. Murata Y, Colonnese MT. GABAergic interneurons excite neonatal hippocampus in vivo. Sci Adv. 2020;6(24):eaba1430.

72. Werner Y, Mass E, Ashok Kumar P, Ulas T, Handler K, Horne A, et al. Cxcr4 distinguishes HSC-derived monocytes from microglia and reveals monocyte immune responses to experimental stroke. Nat Neurosci. 2020;23(3):351–62.

73. Bankhead P, Loughrey MB, Fernandez JA, Dombrowski Y, McArt DG, Dunne PD, et al. QuPath: Open source software for digital pathology image analysis. Sci Rep. 2017;7(1):16878.

74. Hothorn T, Bretz F, Westfall P. Simultaneous inference in general parametric models. Biom J. 2008;50(3):346–63.

75. Bray NL, Pimentel H, Melsted P, Pachter L. Near-optimal probabilistic RNA-seq quantification. Nat Biotechnol. 2016;34(5):525–7.

76. Love MI, Huber W, Anders S. Moderated estimation of fold change and dispersion for RNA-seq data with DESeq2. Genome Biol. 2014;15(12):550.

77. Jin X, Simmons SK, Guo A, Shetty AS, Ko M, Nguyen L, et al. In vivo Perturb-Seq reveals neuronal and glial abnormalities associated with autism risk genes. Science. 2020;370(6520).

78. Hao Y, Stuart T, Kowalski MH, Choudhary S, Hoffman P, Hartman A, et al. Dictionary learning for integrative, multimodal and scalable single-cell analysis. Nat Biotechnol. 2024;42(2):293–304.

79. McGinnis CS, Murrow LM, Gartner ZJ. DoubletFinder: Doublet Detection in Single-Cell RNA Sequencing Data Using Artificial Nearest Neighbors. Cell Syst. 2019;8(4):329–37 e4.

80. Telley L, Agirman G, Prados J, Amberg N, Fievre S, Oberst P, et al. Temporal patterning of apical progenitors and their daughter neurons in the developing neocortex. Science. 2019;364(6440).

81. Andreatta M, Carmona SJ. UCell: Robust and scalable single-cell gene signature scoring. Comput Struct Biotechnol J. 2021;19:3796–8.

82. Park SR, Namkoong S, Friesen L, Cho CS, Zhang ZZ, Chen YC, et al. Single-Cell Transcriptome Analysis of Colon Cancer Cell Response to 5-Fluorouracil-Induced DNA Damage. Cell Rep. 2020;32(8):108077.

83. Finak G, McDavid A, Yajima M, Deng J, Gersuk V, Shalek AK, et al. MAST: a flexible statistical framework for assessing transcriptional changes and characterizing heterogeneity in single-cell RNA sequencing data. Genome Biol. 2015;16:278.

84. Rahmati V, Kirmse K, Holthoff K, Kiebel SJ. Ultra-fast accurate reconstruction of spiking activity from calcium imaging data. J Neurophysiol. 2018;119(5):1863–78.

